# Functional Dissociation in the Developing Superior Temporal Lobe: Language and Theory of Mind

**DOI:** 10.1101/2025.07.25.665611

**Authors:** Kelly J. Hiersche, Laura Bradley, David E. Osher, Zeynep M. Saygin

## Abstract

Language and theory of mind (ToM; the ability to infer others’ mental states) are both crucial for human communication and yet their developmental origins are unclear. Are their neural substrates distinct within the superior temporal lobe (STL) but with opposing lateralization, as in adults? Or do they emerge from common neural substrates during development, perhaps in homologous regions originally involved in more basic social processing? Here we investigate the development of this functional dissociation, and the dissociation of their underlying connectivity fingerprints in a large cohort of children (ages 4-9 years, n=58 sessions) and adults (n=29). We demonstrate for the first time that children show distinct patterns of neural specificity for language and ToM in STL, just like adults. Children show no evidence of developmental ‘disentangling’ cross-sectionally or longitudinally. Finally, children’s connectivity fingerprints predicting future language or ToM activation are almost identical to concurrent fingerprints and are largely non-overlapping across domains. While linguistic and ToM processing undergo continued neural specialization to reach the mature adult-like state, they are distinct remarkably early in human development. Our results challenge the idea that language develops from neural processors common for social communication and instead support distinct neural origins of these mental domains.

## Introduction

Language is a uniquely human skill and vital for effective communication. The neural architecture for linguistic processing is left-lateralized (Malik-Moraleda et al., 2021; Ojemann, 1991), while other types of social communication including theory of mind (ToM; the ability to infer others’ mental states) are right-lateralized in homotopic cortex (Gupta et al., 2025), prompting theories that linguistic neural processors may have emerged in humans from regions originally involved in understanding social environments, a common and vital skill shared across animals (Rajimehr et al., 2022). If true, one prediction would be that language and social processing share common neural architecture especially in homotopic cortices, and perhaps that they may emerge from this common cortex in development as children acquire these skills.

In adults, language and social processing, especially ToM, interact to help us infer what others know and communicate effectively (Fedorenko et al., 2024). However, their neural processors may not overlap. Previous work has directly compared neural responses to language and ToM using a within-subject design, finding that language regions (left hemisphere (LH) temporal regions and frontal regions) have specific, significant univariate responses to language, and not ToM, whereas ToM regions (temporal parietal junction (TPJ), medial prefrontal cortex (mPFC), precuneus) only respond to ToM, not language (Shain et al., 2022). Further, intersubject correlations of subject-specific (for review of importance of subject specific localizers, see (Fedorenko, 2021; Saxe et al., 2006)) language and ToM functional regions suggest modular structures, with no overlap in language and ToM processing in adults (Paunov et al., 2022). Even within the superior temporal lobe (STL) which contains hubs of both language and ToM networks, these two communicative functions are distinct in adults: STL subregions that support language are not selective for ToM and vice versa (Hertrich et al., 2020; Paunov et al., 2019; Shain et al., 2022). Does the same hold true in the developing brain? Given behavioral evidence that language and ToM are intertwined across development (de Villiers, 2021; de Villiers & de Villiers, 2014; Miller, 2006; Slade & Ruffman, 2005), and that both ToM and linguistic skills continue to develop over early childhood (Garfield et al., 2001; Lenneberg, 1967; Perner et al., 1987; Sharp & Hillenbrand, 2008), perhaps the neural processors supporting ToM skills may emerge from common substrates as those that support early linguistic skills.

Brain networks supporting language and ToM are established early (by the age of 3), though they continue to develop over many years (Hiersche et al., 2024; Olulade et al., 2020; Richardson et al., 2018). In developmental studies, these two networks have primarily been examined in isolation but they show similar specificity to adults, with language regions in the frontal and temporal lobe showing selectivity to semantics and syntax as compared to control conditions (Enge et al., 2020; Hiersche et al., 2024; Olulade et al., 2020; Weiss-Croft & Baldeweg, 2015), while ToM regions (regions within STL, temporal parietal junction (TPJ), medial prefrontal cortex (mPFC), and precuneus (PC)) respond more to thinking about others’ mental states (mentalizing; thinking about someone’s beliefs, desires, and emotions) than their bodily sensations, e.g. watching someone experience pain (Richardson et al., 2018; Richardson & Saxe, 2020). However, the development of this neural specialization remains largely unexamined. Are language and ToM represented by spatially and functionally distinct regions within the STL, even in early childhood when ToM skills are still emerging (Rakoczy, 2022) and linguistic skills are still maturing (Oller et al., 2012)? Or are they neurally intertwined, mirroring behavioral evidence, and only becoming more distinct over development, perhaps by strengthening their within network connections and separating from other networks only through repeated co-activations over years of experience (i.e. Interactive Specialization, Johnson, 2011)?

Further, if language and ToM processing are in fact distinct from an early age, what are the neural mechanisms that support their distinct functions? The connectivity fingerprinting hypothesis suggests there is a crucial relationship between a region’s connections and its function (Passingham et al., 2002); comparing differences in connectivity fingerprints can help us understand particular aspects of brain organization (e.g. Mars et al., 2018). Prior work has leveraged this connectivity-function relationship to predict individualized task activation based only on that individual’s connectivity patterns, including for activation to high-level visual categories (Molloy et al., 2024), language (Tavor et al., 2016; Tik et al., 2023), social cognition (Tavor et al., 2016; Tik et al., 2023), and others (Osher et al., 2016, 2019; Saygin et al., 2012; Tobyne et al., 2018). Examining these predictive models reveal the connections that are important for functional specialization, and by comparing models across domains, we can begin to understand the mechanisms of functional dissociation for e.g. language vs. ToM. Further, by examining these relationships across development, we can unravel when and how this dissociation occurs and changes.

Finally, even if language and ToM are functionally and spatially distinct within the same hemisphere, they may be homologues of one another, supported by shared cortical architecture, but lateralized to opposite hemispheres. Prior work examining group level language and social cognition activation found hotspots of language activation along the LH STL spatially align with hotspots of social processing along the RH STL (Rajimehr et al., 2022). If this holds on the individual level, and/or if LH language activation was supported by similar functional connections as RH ToM processing, this would provide additional evidence for theories that linguistic communication, a form of social cognition, may have emerged in cortical regions that were primarily involved in social processing.

In this paper, we build upon prior work in adults demonstrating the functional segregation of language and ToM within the STL, and first attempt to replicate these findings in a new sample of adults and then examine the functional specialization and dissociation of these representations in children ages 4-9 years old. Further, we use longitudinal data to test for evidence of overlap across cognitive domains or developmental progression of functional dissociation. Finally, we examine what functional connectivity fingerprints predict this specialization. We use connectivity fingerprint modeling to determine the relationship between functional connectivity and individualized, voxel-wise functional activation to language and ToM in the STL. We examine if these connectivity patterns are predictive in children and adults, how stable these predictions are over time, and which regions are most important for determining language and ToM specialization within the STL. Through this, we hope to shed light on the emergence of neural specialization of these two domains in humans and the role of connectivity in supporting this specialization.

## Methods

### General Approach

This research builds upon the extensive evidence that language and ToM are functionally and spatially distinct in the mature, adult brain (Braga et al., 2020; Paunov et al., 2019, 2022; Shain et al., 2022). We replicate this dissociation in a new sample of adults, demonstrate similar specificity in a developing sample, and examine the underlying functional connectivity patterns that support language and ToM in the developing and mature state. We specifically focus on STL, as the STL contains specialized regions for both language and ToM, and if there is overlap across these networks, this is where it would be expected (Deen et al., 2015; Paunov et al., 2019). We use functional localizers to compare the functional responses of language defined and ToM-defined regions within the STL. Language and ToM would be considered functionally distinct under two circumstances: 1. Language fROIs are only selective to language (showing no significant mentalizing responses) and ToM fROIs are selective to mentalizing content (showing no significant language responses; see (Shain et al., 2022); 2. Alternatively, we may find that both language and ToM fROIs are significantly responsive to both language and ToM, but they show significantly higher responses to their preferred category (e.g., language responses dwarf those of ToM in language fROIs), as has been demonstrated in prior studies examining e.g. visual category selective regions (see (Kanwisher, 2010; Li et al., 2024)). Further, if language and ToM are primarily spatially distinct, we would expect to see low overlap in their spatial location (i.e., the most language responsive voxels are not the same as the most mentalizing responsive voxels). We also can further examine theories that language and social cognition are homotopes of one another (see (Rajimehr et al., 2022)), by examining overlap of maximally responsive LH language regions and RH ToM regions within individuals. Finally, functionally distinct networks are also separable during rest (Kraus et al., 2021; Smith et al., 2009), and should be supported by distinct patterns of functional connectivity. Therefore, we use functional connectivity to predict the functional responsiveness of each voxel in the STL and compare the important predictors to better understand the neural mechanisms that support the distinction of language and ToM in the STL. If language and ToM are functionally distinct, we also expect a distinct set of predictors (i.e. connections to other regions) to be important for predicting each function.

### Participants

73 children were recruited as part of an ongoing study to examine the relationship between function and connectivity of the developing brain. 50 children completed structural imaging, two runs of an auditory language localizer task, and a ToM task where they watched a short non-verbal movie. 21 of these subjects completed at least one run of both tasks in at least two timepoints (each visit separated by approximately one year, 77 total sessions). Only subjects with a reliable language and ToM in the STL (at least one language functional region of interest (fROI) with positive Sn and Sn>Tx and at least one ToM fROI with positive Mentalizing and Mentalizing>Pain; see Defining fROI section) were included in the cross-sectional analysis (18 sessions excluded; no reliable responses in at least one timepoint). If multiple timepoints had a reliable response, we included all possible sessions. The final cross-sectional sample included 58 sessions across 44 children (mean (sd) age: 6.27 (1.49) years, min-max age: 3.15-9.07 years, 31 female, 44 White/Caucasian, 3 Biracial, 4 Asian, 2 Latino, 5 Black). For any subject in the cross-sectional sample with at least one earlier timepoint available, we selected the most recent earlier timepoint and this became their first timepoint in the longitudinal sample (N=20, mean (sd) between timepoints: 1.09 (0.49) years, 10 female, 5 left-handed, 14 White/Caucasian, 3 Biracial, 2 Black, 1 Latino). Of the cross-sectional subjects, 44 had resting state data with acceptable motion (see Quantifying Motion section), and 15 subjects had an earlier timepoint with low motion resting state data and were therefore included in the longitudinal connectivity fingerprinting analysis.

63 adults were collected as part of an additional study examining the functional organization and connectivity of the mature human brain. 36 adults completed two runs of the language localizer and one run of the ToM localizer (typically in one visit, 6 adults returned for separate scan sessions to collect missing data). Only adults with reliable language and ToM response (same threshold as children) were included in the cross-sectional sample (1 missing language, 6 missing ToM, 7 excluded), leaving a final sample of 29 adults (mean (sd) age: 25.18 (7.77), min-max age: 18.01-55.66, 18 F, 1 left handed, 18 White/Caucasian, 1 Middle Eastern, 4 Asian, 3 Black, 1 Middle Eastern, 1 Latino, 1 Biracial, 1 not provided). As a primary goal of the study is to examine the functional specificity of the language and ToM, we only included subjects with reliable responses to each condition in expected ROIs as above for children.

Both studies were approved by the institutional review board at Ohio State University. Parents gave written consent while children gave verbal assent; adult participants gave written consent. All subjects were compensated monetarily for their time.

### Data acquisition and preprocessing

To make children more comfortable and increase likelihood of collecting high-quality data, many steps were taken prior to and during the scan session. For the child scanning protocol, children arrive approximately one hour prior to the scan. They are familiarized with the study staff, including the staff member running the tasks and monitoring data collection from the control room, and a second member, the ‘buddy’, who stays with the child through the scan. Children are also given the option of keeping a stuffed dog with them for the duration of preparation and scanning. First, children practice the computer ‘games’ (tasks) outside of the scanner to ensure they understand the instructions and to practice keeping their eyes on the screen and head still. Next, they go to a mock scanner, practice lying still and hear the scanner sounds, so that they know what to expect. Finally, they enter the real MRI scanner to begin the scan. Throughout these activities, children fill out a space map with stickers that notes the completion of each preparation step. Foam padding is used during the scan to increase comfort and minimize head motion during the scan. Further, data quality is visually assessed in real-time by the study staff, with the assistance of real-time motion tracking during functional scans. Qualitative assessments of data quality are made by trained study staff; when data quality is low (typically due to motion) scans are either stopped to remind the child to lie as still as possible (e.g., ‘pretend you are frozen like a statue’) or repeated. The scan buddy also monitors the subject to ensure they are lying still, awake, and paying attention to each of the games, and can stop scans if child is confused or needs a break. If needed, children are given a break from the scan to have a snack and move around before continuing the scan. The order of tasks presented are also adjusted to keep the child engaged, though structural scans were always completed first while the child watched a video of their choice to increase their comfort in the scanning environment. Children are compensated with money as well as small toys they picked out as a prize for each “game” they completed. These procedures help ensure that children are comfortable, and the data is of the highest possible quality.

For adults, the same basic quality check procedures and scan order was followed, but they were not familiarized in the mock scanner prior to the scan (unless they preferred to do so), were not given a buddy in the scan room, and were not given breaks during the scan (unless requested).

For both groups, all images were acquired on a Siemens Prisma 3T scanner with a 32-channel phase array receiver head coil and data were analyzed with Freesurfer v.6.0.0, FsFast, FSL, and custom Matlab (version R2020a) code.

#### Anatomical

For both groups, the first scan completed was a whole-head, high resolution T1-weighted magnetization-prepared rapid acquisition with gradient echo (MPRAGE) scan (repetition time (TR) = 2300 ms, echo time (TE) = 2.9ms, voxel resolution = 1.00 mm^3^). Structural data was processed using a semi-automated processing stream (recon-all from FreeSurfer) including intensity correction, skull strip, surface co-registration, spatial smoothing, white matter and subcortical segmentation, and cortical parcellation. The Destrieux atlas cortical parcellation (Destrieux et al., 2010) was used for identifying and masking the superior temporal lobe (combining the superior temporal sulcus and gyrus) on each subject’s brain. Grey matter masks were also created in native anatomy for each subject. Euler number, a number that summarizes the quality of the reconstructed cortical surface file, was calculated for each subject. All subjects had Euler values of 2, which suggests a smooth surface with no holes, and suggests common data quality of structural scans across children and adults. All structural scans, surface reconstruction, and segmentations were also visually examined for defects and subjects were excluded in necessary (one child subject was excluded).

#### Functional

Functional data for localizers and resting-state scans were acquired with multiband 4× accelerated the echo-planar imaging sequence (TR = 1000 ms, TE = 28 ms, voxel resolution = 2 × 2 × 3 mm). For localizer tasks, data were preprocessed with the following steps: motion correction, spatial smoothing (FWHM = 4 mm), and registration to each subject’s anatomical space (using bbregister, (Greve & Fischl, 2009)). Data were further processed in subject’s native anatomy. The resting-state scan was motion corrected and registered to each subject’s native surface space using FreeSurfer’s FS-Fast preprocessing (https://surfer.nmr.mgh.harvard.edu/fswiki/FsFastAnlysisBySteps). Resting state data were further preprocessed using grand median scaling, smoothed on the surface (FWHM = 4mm), linear interpolation over timepoints with greater than 0.5 mm motion (framewise displacement, (Power et al., 2012)), and denoising with nuisance regression for the follow regressors: mean timecourse for white matter and cerebral spinal fluid, top 5 temporal principal components of white matter and cerebral spinal fluid separately, (calculated using CompCor (Behzadi et al., 2007)), and timepoints with greater 0.5mm motion.

ToM Localizer (Jacoby et al., 2016): All participants were instructed to watch the animated movie “Partly Cloudy” (Pixar Animated Studios) once during the scan (and were not familiarized with the movie during scan preparation). Timepoints (346 total) were coded as mentalizing, in which the viewer is thinking about the characters thoughts, or pain, in which the viewer sees the character undergo a physically painful event (see (Jacoby et al., 2016) for more task information and (Richardson et al., 2018) for information on reverse correlation coding). Prior work demonstrates that contrasting these two conditions engages regions of the brain involved in ToM, and has been validated with more traditional ToM tasks, like false belief tasks, but is more accessible for young children (Jacoby et al., 2016; Richardson et al., 2018).

Language localizer (Fedorenko et al., 2010): All participants completed two runs of a language localizer (244 timepoints per run) to identify regions of the brain responsive to auditory language. This task contains blocks of meaningful sentences, nonsense sentences (controlling for prosody but constructed from phonemically intact nonsense words), and texturized speech (controlling for low-level auditory features). Prior work demonstrates that contrasting the sentences with the nonsense conditions engages regions of the brain responsive to the features of high-level language (semantics and syntax). Each run contained 4 blocks (three trials per block, 6 seconds each) of each condition, separated by a 14 second fixation. Each trial ended with a visual queue to press a button, though children who exhibited high motion were instructed to passively listen to the sounds and were not given button boxes, to reduce head motion of trying to look at the button box.

Resting state: During the resting state scan, subjects were instructed to “relax and have a starting contest” with the white crosshair on a black background. During child scans, an experimenter in the scan room watched to see if the child was awake, still, and following instructions. The resting state scan lasted for approximately 5 minutes for all children, and approximately 10 minutes for all adults.

For the longitudinal fROI analysis, data from timepoint two was registered to the timepoint one brain (using mri_robust_register) (Reuter et al., 2010). All registrations were visually examined and passed quality control.

#### Quantifying motion

Mean framewise displacement (Power et al., 2012) and the number of high motion timepoints (greater than 1mm total vector motion, sum of x, y and z dimensions) was calculated for both localizers and the resting state scan. Subjects with greater than 25% high-motion timepoints in either run of the language localizer or the ToM localizer were excluded, and subjects with greater than 25% high-motion timepoints during the resting-state scan were excluded from resting state analyses. Motion across localizers were compared separately for children (for cross-sectional and longitudinal samples) and adults using paired samples t-tests, to ensure that differences between language and ToM activation were not due to motion differences. Motion was initially higher during the language localizer; therefore, one child was removed from the sample to ensure that motion did not significantly differ between language and ToM tasks for the full cross-sectional sample of children (t(55)=1.96, p=0.055). Further, framewise displacement was included as a factor in analyses comparing language and ToM responses. Motion across tasks was not significantly different for the subset used for CF modeling (t(28)=1.05, p=0.30) or adults (t(28)=1.47, p=0.15). Motion was also comparable across timepoints for the fROI longitudinal sample (language localizer: t(19)=1.89, p=0.075; ToM localizer t(19)=1.26, p=0.22) and the CF modeling longitudinal sample (resting state scan: t(14)=1.20, p=0.25). Motion was compared between groups of children and adults but were expected to be significantly different, and children did show significantly greater motion than adults for both tasks (language: t(71)=6.67, p=4.61×10^-9^; ToM: t(71)=6.24, p=2.85×10^-8^). For these analyses, adults were meant as a qualitative benchmark of expected functional segregation, and direct comparisons were not made across child and adult samples, given motion differences. Mean framewise displacement was also included as a covariate for all analyses examining age related changes, to account for typically lower motion in older children.

#### Task fMRI analysis

Preprocessed task data were registered to each subject’s down sampled native anatomical brain (from 1×1×1 mm^3^ to 2×2×2 mm^3^) and included in a volume-based, first-level GLM analysis, with a regressor for each condition of interest (ToM: mentalizing and pain; language: sentences (Sn), nonsense (Ns), texturized (Tx)), as well as motion nuisance regressors (x, y, z, roll, pitch, yaw, high-motion timepoints). The GLM used a block design with a standard boxcar function convolved with the canonical hemodynamic response function (standard gamma function [*d* = 2.25 and *t* = 1.25]). Relevant contrast (ToM: Mntl>Pain; language: Sn>Tx) significance and t-value maps were then resampled to 1×1×1 mm^3^ and used for further analyses including to project on the surface. Given that some analyses require two independent sets of task data, GLMs were analyzed for both runs of the language task separately, as well as combined (for analyses that do not require sets of independent data). Given that children only completed one run of the ToM localizer (due to time constraints), the ToM GLM was run using a full run of data, as well as split-half with the first and second half of timepoints in each condition coded separately (four conditions: Mentalizing 1, Pain 1, Mentalizing 2, Pain 2). For adults with data collected across multiple days, all task data was registered to their first timepoint using mri_vol2vol with linear transform array registration.

#### Defining fROIs and selectivity

To examine the functional specificity of the STL, we created subject-specific functional regions of interest (fROIs) using the group constrained subject specific method (Fedorenko et al., 2010) in three bilateral regions typically activated by high-level language ((Fedorenko et al., 2010); https://evlab.mit.edu/funcloc/; anterior temporal (langAT), posterior superior temporal (langPT), and angular gyrus (langAG)); one bilateral speech region ((Fedorenko et al., 2010); https://evlab.mit.edu/funcloc/; used to ensure language regions did not overlap with speech regions), and three regions typically activated by ToM ((Dufour et al., 2013), https://saxelab.mit.edu/use-our-theory-mind-group-maps/, right superior temporal (r-tomSTS), and left and right temporal parietal junction (tomTPJ)). fROIs were defined for each run (or split half of the data) as the top 5% most significantly active voxels to the contrast of interest (language: Sn>Tx; ToM: Mentalizing>Pain) within a search space and were defined to ensure no overlap in voxel assignment across fROIs (no voxel could be assigned to multiple fROIs). The mean percent signal change (beta estimate multiplied by 100) for each condition was extracted from each fROI in an independent set of data. For example, language and ToM fROIs would be created using the run 1 of the language data and the first half of the ToM data, then PSC would be extracted from run 2 of the language data and the second half of the ToM data. This was then done in the reverse order, and PSC in each fROI was averaged across those two values. This averaged PSC was used to calculate a selectivity value for each fROI to language (Sn – Tx) and ToM (Mentalizing – Pain).

To define longitudinal fROIs, the resulting significance map from the language GLM (combined across run 1 and run 2) and the GLM with the full run of ToM data at the second timepoint was registered to the earlier timepoint brain, and then fROIs were created following the above procedure. Then, the PSC was extracted from the timepoint one data, using a full run of partly cloudy data and the combined language run 1 and 2 GLM (two subjects only had one run of language, and therefore only one run from the other timepoint was used to extract PSC). This allowed us to examine the functional maturation in a similar anatomical region as the later, more mature and selective regions for these fROIs.

#### Defining maximally responsive regions and overlap

Next, we examined spatial overlap in the maximally responsive regions of language and ToM in the STL. The above fROI analysis confined the location of the language and ToM to predetermined search spaces, required us to split the ToM data, and removed any possible overlap in voxels assigned to each fROI; therefore, we also created maximally responsive (hotspot) fROIs using the entire STL (combined superior temporal lateral gyrus and superior temporal sulcal parcels from the Destrieux atlas, (Destrieux et al., 2010)) as a search space, two runs of language, and a full run of the ToM localizer. Overlap (dice coefficient) between most responsive voxels to the contrasts of interest (language: Sn>Tx; ToM: Mentalizing>Pain) was quantified across a series of thresholds (top 1%-5%, 10%, 20%, 30% percent of voxels within the STL). This process was also repeated for each child’s timepoint one data, which allowed us to examine if language and ToM become more spatially distinct across development.

We also tested whether the language hotspots in the LH fall in similar spatial locations to the ToM hotspots in the RH. First, we did a group level analysis to confirm we saw the expected lateralization (language on the LH and ToM on the RH). We computed surface-based laterality index (LI) maps for each participant by projecting individual-level volumetric contrast maps for each task onto each subject’s cortical surface using mri_vol2surf, and then registering them to the symmetric template surface (fsaverage_sym using mris_apply_reg). For both tasks laterality indices were calculated as the difference in activation between the left and right hemispheres (LH – RH). The resulting LI maps were smoothed using an 8 mm full-width at half-maximum (FWHM) Gaussian kernel, and we used a group-level GLM analysis (mri_glmfit) with y cluster-based correction for multiple comparisons via permutation testing (mri_glmfit-sim, 500 permutations, vertex-wise threshold z > 2.0, cluster-wise p < 0.01). The subject-level significance maps on the symmetric template were also used to calculate overlap of language of language hotspots in the LH and ToM in the RH across a series of thresholds (1%-5%, 10%, 20%, 30%).

### Statistical Analyses

First, we examined if the fROIs were selective to their preferred category and functionally distinct (significantly most responsive to their preferred category). To determine if an fROI was significantly selective to either language or ToM we used one-tailed, one-sample t-tests. To examine if these fROIs were also functionally distinct, we used a repeated-measures ANOVA to compare selectivity across tasks. Each rm-ANOVA included a main effect for task and motion (framewise displacement), and subject as the within-subject factor. Following a significant main effect of task, pairwise-tests were used compare selectivity across tasks. A rm-ANOVA was run for the left and right hemisphere language regions separately (l-lang & r-lang; average of anterior and posterior temporal regions within hemisphere) and the left and right hemisphere ToM regions (l-ToM & r-ToM which represented l-tomTPJ and r-tomTPJ). Further, to extend prior work that exclusively examined core language and ToM regions (Shain et al., 2022), we also run a rm-ANOVA for regions on the margins/periphery (Chai et al., 2016; Hertrich et al., 2020; Paunov et al., 2019) of the networks (l-langAG, rlang-AG, r-tomSTS), to determine the extent of functional specificity in the mature (adult) and developing (child) STL. Statistical tests were run separately for children and adults. ANOVAs were run in RStudio (version 2024.12.0+467) using the anova_test function. To examine whether language and ToM disentangle or become progressively more distinct across development, we correlated language and ToM selectivity in each fROI with age in our cross-sectional sample. For example, if selectivity to the non-preferred category decreased with age, that would suggest increasing functional dissociation of language and ToM with age. Selectivity to each task within longitudinal fROIs were compared using paired samples t-tests, to determine if the functional specificity of later matured regions is present earlier in development. Bonferroni-holm multiple comparison correction was used to correct p-values for 7 total comparisons. All statistics, except rm-ANOVA were done in Matlab (R2020a). Across analyses, outliers (+/- 3 standard deviations from the mean) were removed for each fROI (or threshold) separately for all analyses.

Next, we used the hotspot fROIs to determine if language and ToM responses are spatially distinct within the STL. We compared the within hemisphere overlap of language and ToM as well as the across-hemisphere overlap (LH language and RH ToM) at each threshold using two-sample t-tests (and used Bonferroni-Holm correction for 8 threshold comparisons). We compared the overlap in the LH, RH, and across hemispheres using paired-sample t-tests. In the child sample, we correlated overlap with age (controlling for mean framewise displacement across both tasks) to determine if spatial segregation changed across age and used paired-samples t-tests to compare overlap across timepoints, to determine in spatial segregation changed across development.

#### Connectivity Fingerprint Models

Two sets of CF models were created for each subject (for language and ToM). For each model, seed to target functional connectivity (FC), was used to predict functional activation, where the seed was each vertex in the STL, and the targets were all other parcels (based on the (Destrieux et al., 2010) parcellation) in the brain. Models were created separately for each hemisphere and trained separately for child (N=29) and adult (N=29) samples. The first set of models predicted language activation, and the second set predicted mentalizing activation. A separate model was used for each hemisphere.

#### Functional Connectivity

Preprocessed resting state timeseries data was registered to FsAverage symmetric surface space using FreeSurfer’s surfreg and mri_surf2surf. Then, FC was calculated as the correlation between the timeseries in each STL vertex with the mean timeseries across each parcel. FC values were Fischer’s Z transformed within each subject and used as the input for the connectivity fingerprint (CF) models.

#### Task data

The output of the individual subject GLM from the combined two runs of the language task and full run of the ToM task was used in the CF models. The t-statistic maps (language: Sentences>Texturized, ToM: Mentalizing>Pain) for each task were registered to each subject’s surface space with 4mm smoothing, and then to FsAverage symmetric space, following the same procedure as the resting state data. Task data was subset to only include voxels within the bilateral STL parcel. All task data was visually examined on FsAverage symmetric space, along with the STL parcel, to ensure language and ToM hotspots fell primarily within the bounds for all subjects. The functional data was z-scored by subject prior to modeling, and an intercept term was added to each subject’s model.

#### CF Models

We designed *ℒ*_2_ regularized linear ridge regression models, to predict functional activation based on connectivity (**Figure 1**). For each model, the connectivity of each vertex of the STL (9202 per hemi on the FsAverage symmetric brain) to all parcels in the Destrieux Altas (72 per hemisphere and the contralateral STL, 145 total) predicted either the language selectivity, ToM selectivity, or language-ToM selectivity. A separate model was created for each subject, and to avoid overfitting, a nested cross-validation approach was used. Leave-one-out “outer loops” ensured the testing subject was independent from all model training. Leave-one-out “inner loops” were used for optimizing the λ regularization hyperparameter (100 potential values defined as a logarithmically spaced vector of values between 10^-5^ and 10^2^). In each loop, all of the data of a left-out subject were excluded from model design and then used to assess model performance. The optimal λ was selected as the value that maximized the average mean squared error of the held-out inner-loop subject for each possible λ across all inner loops. Then, the mean model coefficients (beta weights) across all inner loops for that optimized λ were computed, and the resulting vector was multiplied by the connectivity matrix of the outer loop left-out test subject, to predict that subject’s functional activation. The predicted functional activation was correlated with the true functional activation, providing a model performance (predicted correlation) value for each hemisphere of each subject, for each model set (language and ToM).

**Figure 1:**
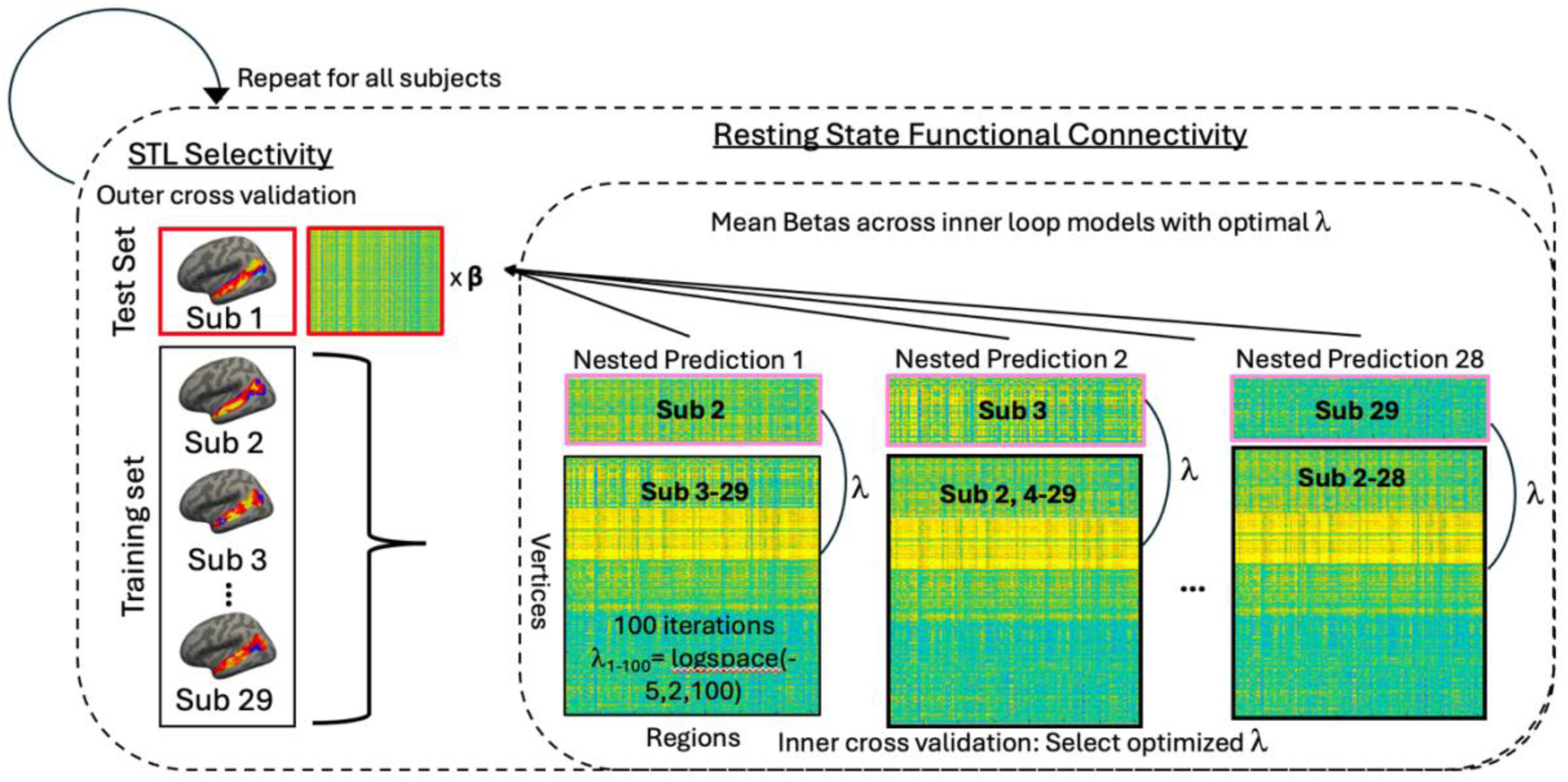
Model fitting schematic. For children and adults separately, we trained a model to predict functional selectivity in the STL (language, ToM), using that same subject’s functional connectivity at rest (each vertex connectivity to all other parcels in the brain). A separate model was constructed for each hemisphere. A nested cross validation loop was used to optimize the λ hyperparameter used for ridge regression, without overfitting the model. The held out outer subject is highlighted in red, whereas the held-out subject for each inner-loop is noted in pink.

#### Evaluating model performance

Model performance was characterized according to model fit, the correlation (R value) of the predicted voxelwise functional activation and actual functional activation. To determine whether the connectivity models *captured individual variability* in connectivity and function, we compared models using the test subject’s own connectivity data vs. another subject’s data (self vs. other). A two-tailed, paired t-test was used to compare the model performance between self and mean model performance from all others. Given that model performance values were correlations, all model performances were Fischer’s Z transformed prior to any statistics. As a quality check, model performance was also correlated with mean framewise displacement during the resting state scan, to ensure model performance was not related to motion. Indeed, motion during the resting-state scan was *not* related to model performance for any of the connectivity fingerprinting models (adults’ language: LH: r=-0.038, p=0.85; RH: r=-0.054, p=0.79; adults’ ToM: LH: r=0.13, p=0.52, RH: r=-0.035, p=0.86; children language: LH: r=-0.050, p=0.80, RH: r=0.047, p=0.81; children ToM: LH: r=-0.14, p=0.45; RH=0.15, p=0.43). However, motion was significantly higher in children compared to adults during the resting-state scan (t(56)=5.65, p=5.69×10^-7^).

#### Examining development of the connectivity function relationship

In the cross-sectional sample of children (N=29), we correlated model performance with age. Further, for children with an earlier timepoint available, we applied the beta weights from the timepoint 2 model to their timepoint 1 connectivity data to yield longitudinal predictions per subject, and this predicted functional activation was correlated with the true (actual) timepoint 2 functional activation. We compared (two-tailed paired samples t-tests) the performance of a subject’s connectivity data collected during the same timepoint with the performance when using connectivity data collected one year prior. This analysis allowed us to examine how stable the connectivity-function relationship was within individuals across time.

#### Comparing across models

Model performance was compared across the language and ToM models for children and adults separately, using two-tailed, paired t-tests. Further, given expected lateralization of language activation, model performance across hemispheres was compared for all models using two-tailed, paired t-tests.

#### Evaluating the principal connectivity fingerprint predictors

Beta weights for each parcel were used to determine value of each predictor (brain region) to model. First, to determine if the same connections were important for predicting function across hemispheres, the average beta weights across all subjects’ LH models (within child and adult group) were correlated (Pearson correlation) with the average RH model beta weights for the language and ToM models. Further, to determine if the same predictors were important across model sets and groups, the beta weights for each hemisphere (LH and RH) of each group (kids and adults) for each model set (language, ToM) were correlated with all others, and Bonferroni-Holm corrected (28 total correlations). Further, the top ten most important positive and negative predictors (based on beta weight), for each model set were plotted on the brain and qualitatively compared across groups and hemispheres to examine overlap in important predictors across models, as well as to interpret important connections that drive the specialization of the STL. Finally, to examine differences in important predictors across the language and ToM models, a paired-samples t-test was run to compare the language and ToM beta weights across individuals (within group) for all 145 brain regions, and Bonferroni-Holm multiple corrected across all regions.

### Data and Code Availability

Data used for statistical analyses and plots, as well as analyses code will be made available upon publication here: https://github.com/SayginLab/STL_languageToM.

## Results

### How functionally distinct are language and ToM in the adult superior temporal lobe? Replication of previous work

To examine functional specificity of the language and ToM networks, we defined fROIs for each (language using Sn>Tx contrast; ToM using Mntl>Pain contrast) and examined their selectivity to their preferred and non-preferred stimuli. Aligning with prior work, adults show expected selectivity to language in bilateral language regions; and selectivity to mentalizing in bilateral ToM regions (**Figure 2A; Table S1**). Adults also show activation to the unexpected category for core language and ToM regions (l-lang, r-lang show significant ToM selectivity and l-ToM, r-ToM show significant language selectivity). Importantly, however, all language regions show significantly higher selectivity to language than mentalizing (all p<0.001, **Table 1**), whereas ToM regions show significantly higher selectivity to mentalizing than to language (all p<0.001). Regions on the margins of the language (l-langAG) and ToM (r-tomSTS) networks show undifferentiated, significant selectivity to both language and ToM (all main effect of task p>0.3).

**Figure 2:**
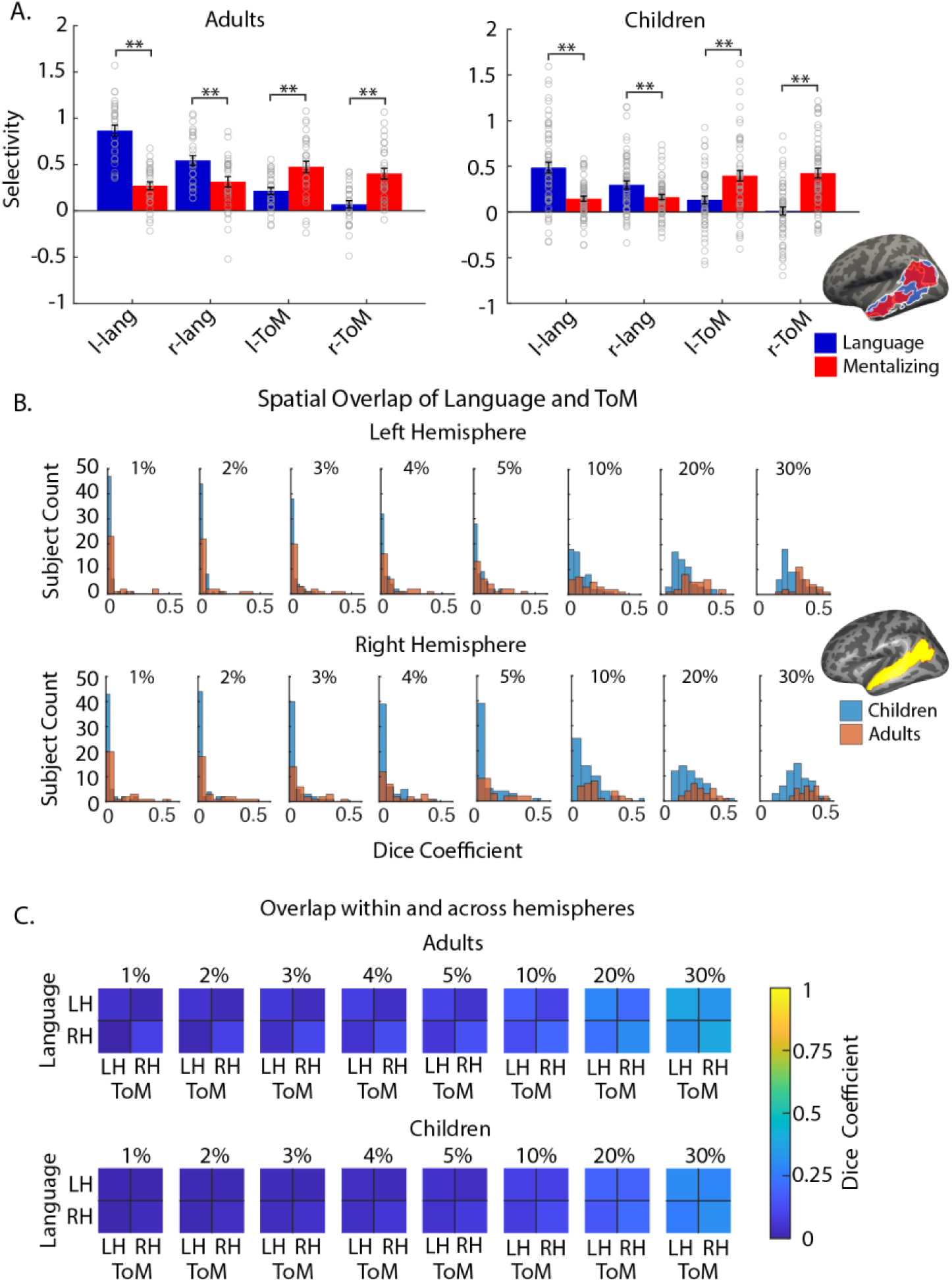
A. Selectivity of language and ToM fROIs. (a) Selectivity for both language (blue) and mentalizing (red) in N=29 adults (left), N=56 children (right), *p<0.05 main effect of task in rmANOVA. Inset demonstrates all language and ToM parcels used in fROI analysis on the LH inflated surface of FsAverage. B. Histograms demonstarting the amount of spatial overlap in maximally sensitive language and ToM regions in the STL across a series of thresholds (LH on top, RH on bottom). Child (blue) and adult (red) histograms are overlayed. Inset shows the anatomical search space used when determining overlap of language and ToM hotspot fROIs. C. Heatmap demonstrating the mean overlap across language and ToM responses within and across hemispheres (adults top, children bottom).

**Table 1:** Results of repeated measures ANCOVA, with main effect of task and framewise displacement included as a covariate, with generalized η^2^ effect sizes listed. Post-hoc t-tests were run to compare selectivity across tasks only after significant effect of task with Cohen’s d effect sizes and Bonferroni-Holm multiple comparison correction (*p<0.05 main effect; **p<0.05 Bonferroni-holm for t-tests to parse main effect of task).

| Adults |  | rmANCOVA results |  |  | post-hoc t-test |  |  |
| --- | --- | --- | --- | --- | --- | --- | --- |
| *fROI* | | *F* | *p* | $\eta^2$ | *t(df)* | *p* | *d* |
| **l-lang** | motion | F(1,55)=0.44 | 0.51 | 0.008 |  |  |  |
|  | task | F(1,55)=68.42 | 3.15x10-11** | 0.55 | t(28)=12.8 | 3.36x10-13** | 2.37 |
| **r-lang** | motion | F(1,55)=0.17 | 0.68 | 0.003 |  |  |  |
|  | task | F(1,55)=9.85 | 0.003** | 0.15 | t(28)=4.47 | 1.17x10-4** | 0.83 |
| **l-ToM** | motion | F(1,53)=0.48 | 0.49 | 0.009 |  |  |  |
|  | task | F(1,53)=14.99 | 0.0003** | 0.22 | t(27)=-4.44 | 1.38e- 4** | -0.8 |
| **r-ToM** | motion | F(1,55)=1.16 | 0.28 | 0.021 |  |  |  |
|  | task | F(1,55)=22.01 | 1.84x10-5** | 0.29 | t(28)=-5.90 | 2.38e- 6** | -1.1 |
| **l-langAG** | motion | F(1,55)=2.63 | 0.11 | 0.046 |  |  |  |
|  | task | F(1,55)=0.093 | 0.76 | 0.002 |  |  |  |
| **r-langAG** | motion | F(1,51)=8.78 | 0.005** | 0.15 | t(28)=1.47 | 0.15 | 0.24 |
|  | task | F(1,51)=0.99 | 0.32 | 0.019 |  |  |  |
| **r-tomSTS** | motion | F(1,55)=0.11 | 0.74 | 0.002 |  |  |  |
|  | task | F(1,55)=0.79 | 0.38 | 0.014 |  |  |  |
| Children |  | rmANCOVA results |  |  | post-hoc t-test |  |  |
| *fROI* | | *F* | *p* | $\eta^2$ | *t(df)* | *p* | *d* |
| **l-lang** | motion | F(1,107)=3.397 | 0.068 | 0.031 |  |  |  |
|  | task | F(1,107)=33.32 | 7.71x10-8** | 0.24 | t(54)=5.83 | 3.18x10-7 | 0.79 |
| **r-lang** | motion | F(1,107)=0.32 | 0.57 | 0.003 |  |  |  |
|  | task | F(1,107)=8.30 | 0.005** | 0.072 | t(54)=2.97 | 0.004 | 0.4 |
| **l-ToM** | motion | F(1,103)=0.19 | 0.67 | 0.002 |  |  |  |
|  | task | F(1,103)=12.74 | 0.00055** | 0.11 | t(52)=-3.26 | 0.002 | -0.5 |
| **r-ToM** | motion | F(1,101)=0.85 | 0.77 | 0.00084 |  |  |  |
|  | task | F(1,101)=31.64 | 1.66x10-7** | 0.24 | t(51)=-5.53 | 1.09x10-6 | -0.8 |
| **l-langAG** | motion | F(1,99)=0.14 | 0.71 | 0.001 |  |  |  |
|  | task | F(1,99)=0.44 | 0.51 | 0.004 |  |  |  |
| **r-langAG** | motion | F(1,91)=0.14 | 0.71 | 0.001 |  |  |  |
|  | task | F(1,91)=0.081 | 0.78 | 0.00089 |  |  |  |
| **r-tomSTS** | motion | F(1,101)=0.75 | 0.39 | 0.007 |  |  |  |
|  | task | F(1,101)=0.00069 | 0.98 | 6.8x10-6 |  |  |  |

### Are language and ToM distinct in the *developing* superior temporal lobe?

Children show a similar pattern to adults, with selectivity for language in the core language regions (all p<0.0001), and selectivity for mentalizing in core ToM regions (all p<0.0001) (**Table S1**, **Figure 2A**). Like adults, children also show selectivity to the non-preferred domain across both language and ToM fROIs. Importantly, and aligning with adults, children’s language and ToM fROIs show distinctive functional selectivity: language regions are significantly more selective to language than ToM (all p<0.01, **Table 1**), and ToM regions are significantly more selective to ToM than language (all p<0.01). And again, as observed in adults, children show no differentiation between the two domains in the margins of the language and ToM networks; r-tomSTS and bilateral langAG are selective to both language and ToM, showing no distinction between them (all main effect of task p>0.5).

To examine whether language and ToM disentangle or become progressively more distinct across development, we correlated language and ToM selectivity in each fROI with age. If language and ToM are disentangling over time, we might expect ToM selectivity in language fROIs to decrease (or language selectivity in ToM fROIs to decrease). However, this was not the case in our sample, as we did not see evidence of selectivity changes to the non-preferred category across age (p>0.05, see **Table S2**). Given that differentiation of language and ToM may come from increased experience and skills which may vary across individuals, we used the longitudinal sample, for whom we would expect skills to improve within subject, to compare language and mentalizing selectivity across timepoints (by projecting their TP2 fROI back to TP1 and extracting selectivity). We observed no evidence of change in selectivity to the non-preferred category over development (all p>0.05, see **Table S2**).

### How spatially distinct are language and ToM in the adult superior temporal lobe?

Next, we examined the overlap in the maximally selective voxels to each task across the STL. This allowed us to examine any spatial overlap in hotspots of activation, as well as overcome potential limitations of the prior analysis (the use of predefined search spaces and different amounts of data for the language and ToM localizers). The most selective voxels along the entire STL (anatomically defined in each subject’s brain) were selected at a series of thresholds for each task separately and then a dice coefficient was calculated to quantify any overlap (1 is complete overlap, 0 is no overlap). Adults show minimal overlap between these ‘hotspots’ of activation at all thresholds (**Figure 2B**), though unsurprisingly, there is higher overlap for the higher (more liberal) thresholds (mean ± sd overlap: LH: 1%: 0.048 ± 0.11, 30%: 0.38 ± 0.092; RH: 1%: 0.090 ± 0.14; 30%: 0.39 ± 0.078; see **Table S3** for all thresholds). Further, given prior work (e.g., Rajimehr et al., 2022) suggesting that while language and ToM may not overlap within hemisphere, they may be homotopes and therefore overlap in the opposite hemisphere (processing of social cognition would develop in the RH homotopes of LH dominant language regions for example), we also compared overlap of LH language responses with RH ToM responses (using a symmetric template brain). Once again, we see minimal overlap across all thresholds (mean ± sd overlap:1%: 0.0037 ± 0.17; 2%: 0.019 ± 0.053; 3%: 0.041 ± 0.078; 4%: 0.058 ± 0.094; 5%: 0.069 ± 0.10; 10%: 0.13 ± 0.13; 20%: 0.23 ± 0.11, 20%: 0.32 ± 0.85), despite seeing expected lateralization of language to the LH and ToM to the RH (see **Figure S1**).

### Are language and ToM spatially distinct in the *developing* superior temporal lobe?

Like adults, children show minimal overlap between ‘hotspots’ of activation across all thresholds (**Figure 2B**), again with expectedly higher overlap in higher thresholds (mean ± sd overlap: LH: 1%: 0.014 ± 0.034; 30%: 0.30 ± 0.088; RH: 1%: 0.030 ± 0.060; 30%: 0.31 ± 0.093, see **Table S3** for all thresholds). Further, there was no evidence of language and ToM disentangling across development: the amount of overlap was not correlated with age at any threshold (all p>0.3; **Table S4**, nor did overlap change across timepoints (all Bonferroni-Holm p>0.2, **Table S4**). Further aligning with adults, children show very minimal overlap of LH language and RH ToM hotspots across all thresholds (mean ± sd overlap: 1% 0.0082 ± 0.041; 2%: 0.017 ± 0.058; 3%: 0.032 ± 0.071; 4%: 0.043 ± 0.085; 5%: 0.052 ± 0.096; 10%: 0.080 ± 0.11; 20%: 0.16 ± 0.11; 30%: 0.25 ± 0.13), with no evidence of changing across ages (all p>0.05, **Table S4**) or development (all p>0.5, Table S4).

### Connectivity fingerprints that underlie STL function

We then explored whether there existed similarities in the underlying connectivity that predicts (and may therefore determine) functional specialization for language and ToM within the STL (i.e. connectivity fingerprints for function). It is possible that despite observing little to no overlap between language and ToM specialization in either adults or children, that the two domains emerge from common spatial locales or connections earlier in development, and that underlying connectivity patterns would reveal these vestigial commonalities at age 4 to adulthood. We first evaluated whether connectivity successfully predicted functional activation to ToM and language in both adults and children, and then we explored the similarities and differences in the connectivity fingerprints for ToM vs. language in both cohorts.

#### Predicting Adult STL Activation

Language selectivity was successfully predicted using functional connectivity (predicted correlation: mean ± sd: LH: 0.60 ± 0.24, RH: 0.50 ± 0.22) and a subject’s own connectivity outperformed using another’s connectivity (predicted correlation using other: mean ± sd: LH: 0.36 ± 0.11; RH: 0.34 ± 0.093; self vs. other t-test: LH: t(28)=5.22, p=1.51×10^-5^; RH: t(28)=4.67, p=6.80×10^-5^; see **Figure 3A**), suggesting a subject-specific relationship between connectivity and language activation. This was also true for ToM; selectivity was successfully predicted using functional connectivity (predicted correlation: mean ± sd LH: 0.63 ± 0.25, RH: 0.56 ± 0.27), and a subject’s own connectivity outperformed using another’s connectivity (predicted correlation using other: mean ± sd: LH: 0.24 ± 0.14, RH: 0.22 ± 0.13; self vs other t-test: LH: (t(28)=12.58, p=4.84×10^-13^, RH: t(28)=9.03, p=8.75×10^-10^, see **Figure 3A**).

**Figure 3:**
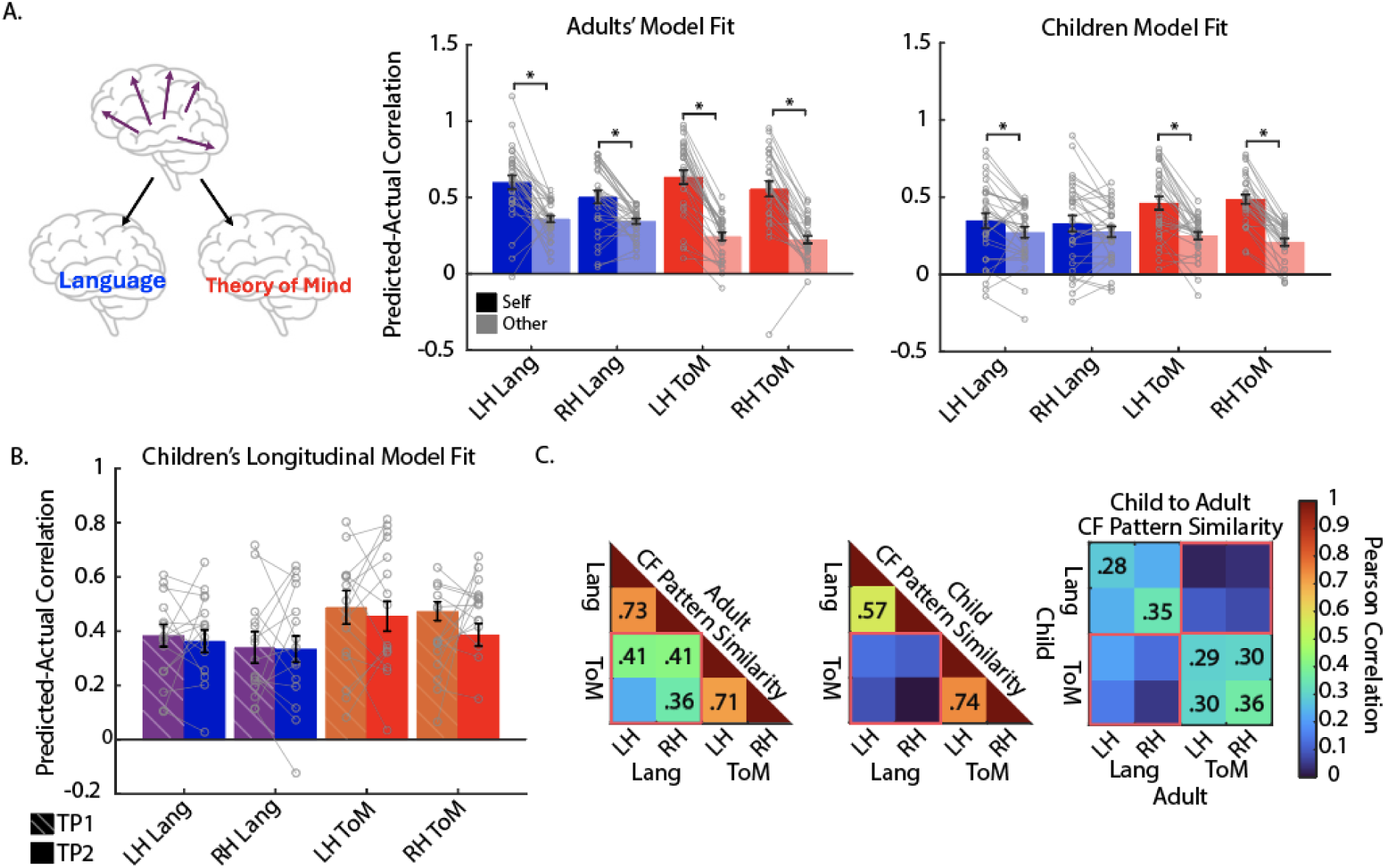
CF model performance. A. Left, schematic of using connectivity of STL to predict function. Center and right, mean Fischer’s Z transform of predicted-actual correlation with standard error, for children and adults respectively. B. Mean Fischer’s Z transform of predicted-actual correlation with standard error bars when using concurrent resting-state data to predict task activation (TP2 connectivity to predict TP2 function) or using data from timepoint 1 (N=15 subjects with longitudinal). Hollow circles represent individual subject model performance. Grey lines connect each subject’s model performance. *p<0.05 two-tailed, paired *t* test. C. Connectivity fingerprint pattern similarity across hemispheres and tasks. Pearson correlation of average (across individuals) beta weights to compare across tasks within adults (left), across tasks within children (middle), and across tasks and groups (right). Significant correlations after Bonferroni-Holm correction for 28 comparisons have their Pearson’s R value listed. Language and ToM comparisons are highlighted in red.

#### Predicting Developing Language Activation

In children, language selectivity was also successfully predicted using functional connectivity (predicted correlation mean ± sd: LH: 0.35 ± 0.28; RH: 0.33 ± 0.28), and a child’s own connectivity outperformed predictions using another’s connectivity but only in the LH (predicted correlation: mean ± sd: LH: 0.27 ± 0.19; self-other t-test: t(28)=2.18, p=0.038), not in the RH (predicted correlation: mean ± sd: 0.28 ± 0.19; self-other t-test: t(28)=1.61, p=0.12, see **Figure 3A**), suggesting a strong relationship between connectivity and function only in the language-dominant LH. This relationship increased with age only in the language dominant LH (correlation of predicted and age controlling for motion: LH: r=0.42, p=0.023; RH: r=0.25, p=0.21).

Next, using our longitudinal data, we examined how stable the connectivity-function relationship is within subject across time. We find that a child’s connectivity data predicted their functional activation to language one year later (TP1-predicted correlation mean ± sd: LH: 0.36 ± 0.16; RH: 0.33 ± 0.19). This baseline connectivity data predicted their later functional activation just as well as their concurrent connectivity data for both hemispheres (t-test between predictions made using TP1 connectivity data vs. TP2 connectivity: LH: t(14)=-0.46, p=0.65, RH: t(14)=-0.10, p=0.92), suggesting a subject-specific, stable relationship across time between connectivity and language activation (see **Figure 3B**).

#### Predicting Developing ToM Activation

In children, ToM selectivity was successfully predicted using functional connectivity (mean ± sd: LH: 0.46 ± 0.23; RH: 0.49 ± 0.16), and ToM activation predicted by a subject’s own connectivity outperformed using another’s connectivity in both hemispheres (mean ± sd: LH: 0.25 ± 0.13; RH: 0.21 ± 0.13; self vs other t-test: LH: (t(28)=5.86, p=2.65×10^-06^, RH: t(28)=10.62, p=2.49×10^-11^, see **Figure 3A**), suggesting a subject-specific relationship bilaterally in the STL. This relationship increased with age but only in the non-dominant LH (correlation of predicted and age controlling for motion: LH: r=0.40, p=0.032; RH: r=0.26, p=0.18).

Again, we examined the stability of the connectivity-function relationship across time, finding that a child’s connectivity data predicted their functional activation to mentalizing one year later (TP1-predicted correlation mean ± sd: LH: 0.46 ± 0.21; RH: 0.39 ± 0.16). Similar to language activation, a subject’s baseline connectivity data (TP1) did as well as their concurrent connectivity (TP2) when predicting ToM activation (t-test between TP1 and TP2: LH: t(14)=-0.62, p=0.54, RH: t(11)=-1.59, p=0.13; see **Figure 3B**).

#### Comparing Across Models for predicting Language and ToM

For adults, language and ToM activation were equally well-predicted in both hemispheres (t-test language vs ToM predicted-actual correlation: LH: t(28)=-0.52, p=0.60; RH: t(28)=-0.90, p=0.37). For children, while ToM and language activation were equally well predicted from connectivity in the LH: t(28)=-1.81 p=0.081), ToM activation was better predicted than language activation in the RH (language vs ToM t-test; t(28)=-2.77, p=0.0097), which is not surprising given that the models failed to characterize a subject-specific relationship between connectivity and language activation in the RH STL.

#### Connectivity fingerprints for Language vs. ToM

Given that both language and ToM were successfully predicted using functional connectivity, we next evaluated the model coefficients (i.e. connectivity fingerprints) to determine which connections are important for predicting an STL voxel’s language or ToM selectivity. To examine potential similarities across models, we correlated the beta weights for the language and ToM models (F**igure 3C**). We found strong pattern similarity within task across hemispheres for both adults (language LH-RH: r=0.73, p_bh_=3.58×10^-24^; ToM LH-RH: r=0.71, p_bh_=2.06×10^-22^) and children (language LH-RH: r=0.57, pbh=2.08×10^-12^; ToM LH-RH: r=0.74, p_bh_=3.55×10^-25^). Child and adult models also showed weaker, but still significant, similarity within tasks (adult-child language: LH: r=0.27, p_bh_=0.011; RH: r=0.35, p_bh_=4,18×10_-4_; ToM LH: r=0.30, p_bh_=0.0052; RH: r=0.36, p_bh_=1.47×10^-4^). Importantly, we examined whether the fingerprints were similar across tasks within hemispheres. Adults do in fact have similar connectivity fingerprints for the language and ToM tasks (LH Lang-LH ToM r=0.37; RH Lang-RH ToM r=0.26); however, children, do not (all r<0.1 and all p>0.05). When we look across tasks and hemispheres, we find that adults again show similarities (LH Lang-RH ToM r=0.36; RH Lang-LH ToM: r=0.19;), though they are much lower than the within task pattern similarity, whereas children do not show similarities across tasks (all r<0.3 and all p>0.05). This suggests that language and ToM do not emerge from similar connectivity fingerprints; however, they may become more similar through development and exposure.

What contributes to these differences across domains? When we compared beta values across language and ToM for each set of models, we found that over 90% of the 145 predictors (connectivity targets) significantly differ across the models (percentage of targets that significantly differ between language and ToM models after Bonferroni-holm correction: adults: LH: 90.34%; RH: 91.03%; children: LH: 97.24%; RH: 91.03%). To make these findings more interpretable, we plot only the top regions that differ across models (for adults see **Figure S2;** for children see **Figure 4**). These qualitative comparisons suggest that connectivity with other nodes of the ToM or language networks are important for predicting each domain respectively. For example, in adults (see **Figure S2**) we see that LH inferior frontal regions are positive predictors for language and negative for ToM (greater connectivity to LH inferior frontal predicts language activation and vice versa for ToM activation), whereas the bilateral angular gyrus and LH medial subparietal regions (precuneus) are negative predictors for language and positive for ToM. Similarly in children (see **Figure 4**), LH inferior frontal, LH motor, and RH insular are positive predictors for language and negative for ToM, whereas the bilateral medial subparietal regions, the RH TPJ, and the precuneus are positive for ToM and negative for language. Further, we plotted the regions that most differ across child and adult models for both language and ToM (see **Figure S3**), and found that some regions show magnitude differences, becoming even more important predictors in adults (e.g., the contralateral STL when predicting ToM, LH inferior frontal regions when predicting language), perhaps reflecting the continued functional and connectional development of these regions. However, we also observe differences in sign for child vs. adult models of less predictive connections (i.e. connections with lower betas), such that for the RH language model, RH medial regions are weakly positive predictors in children, but negative predictors in adults, whereas LH inferior and transverse temporal regions are positive predictors in children, but negative in adults.

**Figure 4:**
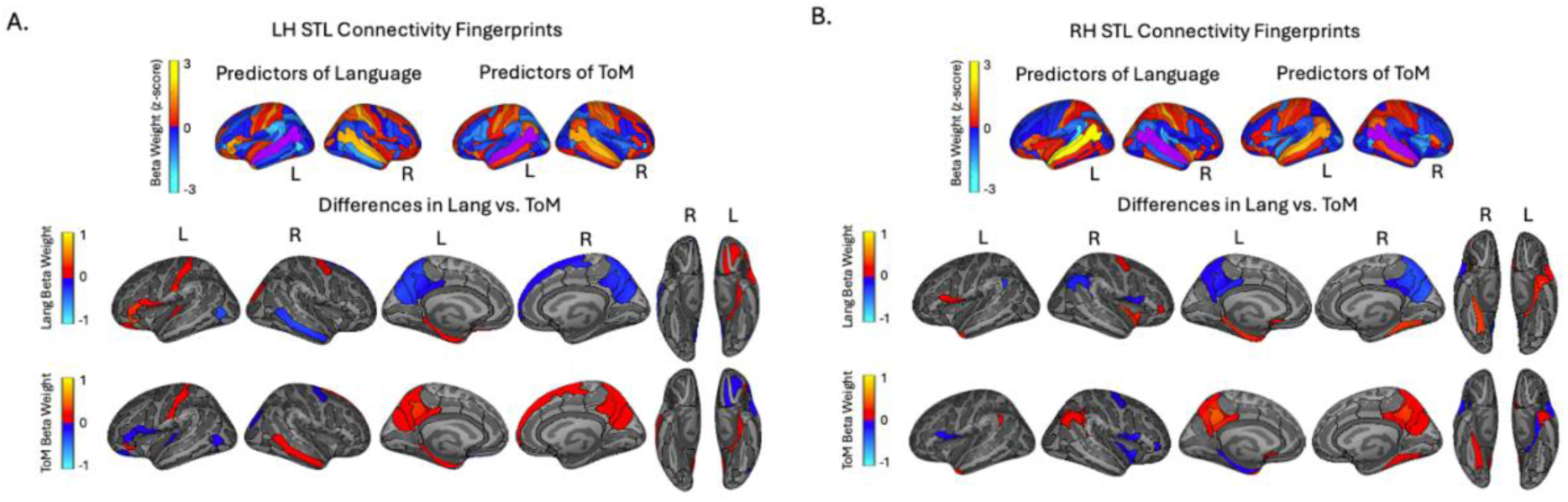
Beta value for the child language and ToM models. Z-score of beta weights for language and ToM models for A. LH and B. RH. Top row shows all beta weights on the lateral surface. The region being predicted is depicted in purple. Below, raw beta weights for regions that are most different when predicting language and ToM (10 most positive and 10 most negative t-values after paired-sample t-tests across language and ToM models) for LH and RH separately. Middle row depicts beta value in the model predicting language activation, bottom row depicts beta value for the model predicting ToM activation.

## Discussion

Here, we ask if language and ToM processing are spatially and functionally distinct in the developing human brain and examine the neural mechanisms that support their differentiation. Prominent theories have suggested that language processing may have emerged in humans from common neural mechanisms as other social communication, such as ToM, and so it is plausible that the two processes are linked (or perhaps occupy homotopic cortices on either hemisphere) early in development and disentangle over time as these skills develop through early childhood. We scanned children as young as 4 years of age on two separate fMRI tasks in order to define subject-specific activation profiles and compare overlap between domains. This study offered a unique opportunity to examine the functional dissociation of language and ToM processing in children whose linguistic and ToM skills are still developing; additionally, we were able to investigate both cross-sectional and within-subject development of this dissociation. Finally, we used resting-state connectivity scans in these children to predict individualized functional activation to either domain and compared these fingerprints across domain, across hemisphere, and across age within individual.

For the first time, we examine the functional specificity of two skills vital for communication, verbal language and mentalizing, during development in young children who are still refining these skills. A key developmental hypothesis of ToM suggests that language development (perhaps generally or syntax development specifically) is fundamental for ToM development (Astington & Jenkins, 1999; de Villiers & de Villiers, 2014; Ebert, 2020; Garfield et al., 2001; Hale & Tager-Flusberg, 2003; Miller, 2006). Other work suggests that ToM and language have similar cognitive building blocks, such as recognizing intention, shared attention, and/or working memory (Baldwin & Moses, 2001; Miller, 2006; Slade & Ruffman, 2005). Further, language and ToM skills are predictive of one another early in development (de Villiers & Pyers, 2002). Given the overlapping nature of these skills behaviorally, it is plausible that they may overlap in the brain; however, this is not what we find. We find that children show functional and spatial differentiation of language and ToM; core language regions are primarily selective to language and core ToM regions primarily selective for mentalizing. Further, we find that language and ToM are already dissociated by age 4 even though these skills are still emerging and developing. These results align with prior work using EEG demonstrating false belief understanding and complementation syntax have distinct neural signatures in elementary school children (Guan et al., 2020), as well as lesion work and studies examining atypical development, both of which suggest that language and social cognition can be impaired independent of one another (Dronkers et al., 1998; Varley et al., 2001; Willems et al., 2011). Prior work has also demonstrated that ToM networks are distinct from others like the pain network, even in young children that do not pass false belief tasks (Richardson et al., 2018). Finally, in addition to the functional independence of temporal language regions demonstrated here, previous work suggests that frontal language regions also do not emerge from more generalized processors, and that these regions are spatially and connectionally distinct from adjacent regions engaged in mental effort and other types of thought (Hiersche et al., 2024; Schettini et al., 2023).

Why might the neural dissociation of language and ToM precede behavioral maturation? If a region’s innate connectivity patterns scaffold its later functional specialization (i.e., connectivity hypothesis), then it is plausible that neural dissociation may precede skill development. Additionally, while we do not see spatial overlap in language or ToM responses functionally within or across hemisphere, these functions may still be linked in their common connections to the rest of the brain and may emerge through shared connectivity scaffolds. Language is a tool for social communication (Tylén et al., 2010), and as such, it may have evolved in LH regions that previously specialized for other social cognitive skills, prior to language emergence (Rajimehr et al., 2022). Therefore, language and ToM may have evolved as homologous substrates in the LH and RH STL. Both skills may rely on common connectivity patterns to initially set them up to process social cues important for effective communication, then become more specialized through ongoing experience-dependent changes in connectivity (as suggested by the interactive specialization hypothesis (Johnson, 2011)). However, this is not what our results suggest. Instead, we found that language and ToM activation are predicted by distinct patterns of connectivity even in children. Similar to adults, we find that a child’s STL connectivity to other language regions (e.g., frontal regions) are important for predicting language responses, whereas connectivity to other ToM regions (e.g., precuneus) are important for predicting higher ToM responses. Additionally, we found little similarity between connectivity fingerprints of domains within hemispheres, particularly in children; in fact, fingerprints were more similar for predicting the same domain across hemisphere than a different domain across or even within hemisphere (i.e. fingerprints for RH language vs. LH language were more similar than RH language vs. LH ToM or RH ToM) suggesting little evidence of vestigial homologies. Given that adults show greater similarity in language and ToM connectivity fingerprints and children show no significant similarities, we do not see evidence that language and ToM emerged from similar connections, but instead that these functions become more intertwined after years of experience and maturation.

Still, it is possible that these processes emerged from a common neural substrate earlier than age 4. Accordingly, we may expect to observe ongoing disentanglement between these domains especially since ToM skills typically emerge between years 4-5 and are likely still developing in our cohort. However, we found that these processes do not seem to further disentangle across development. Functional activation to the nonpreferred domain is stable across ages and timepoints, as is the amount of spatial overlap between language and ToM. Further, our longitudinal predictive analyses suggest that connectivity fingerprints are not different across time within individual, suggesting stable subject-specific relationships between connectivity and function for both language and ToM. Notably however, our between-group comparisons of adult vs. child connectivity fingerprints revealed some developmental differences. While the same predictors are important for child and adult models, the magnitude of their importance in predicting language or ToM increases in adult models. For example, while LH frontal regions positively predict language activation in both child and adult models, these regions are more important (i.e. beta weight is higher) for adults than children. And while the most important language predictor across models for children and adults is the contralateral STL, this cross-hemispheric connectivity becomes even more important in adults, making it also one of the regions with the greatest differences between the child and adult models. This result suggests that crosstalk between the STL across hemispheres becomes increasing important across development. Altogether, these results suggest that while connectivity fingerprints are relatively stable within subject in early childhood, there are certainly some developmental changes that occur after age 9. These changes likely reflect ongoing maturational changes within domain but do not appear to reflect further dissociation between ToM and language processes.

In order to understand these processes in young children, we used simple tasks (movie-watching and listening) to overcome previous obstacles in developmental data collection. However, ToM is just one aspect of social cognition that has been implicated in the STL, and various other social cognitive and communicative skills (e.g., eye gaze, joint attention) are associated with behavioral development of language and ToM (Baldwin & Moses, 2001; Freiwald, 2020; Miller, 2006). Therefore, it would be interesting for future studies to examine the development and neural specificity of additional social cognitive building blocks, with respect to ToM and language. Skills such as joint attention may be particularly important to examine, given that it is one of the building blocks of ToM and word learning in young children (Slade & Ruffman, 2005). Further, future work should explore whether ongoing separation and refinement of connectivity patterns determine ongoing functional specialization. Additionally, while our sample includes the youngest possible ages of children who can be tested on an extensive battery of structural and functional imaging, it is possible that we may find overlapping activation or CF patterns in younger children (perhaps relying more on unimodal sensory cortices that develop earlier (Skeide & Friederici, 2016). Future work should also examine the CFs that drive atypical functional organization of language and ToM (e.g., under case of language remapping to the RH (follow hemispherectomy, (Ivanova et al., 2017; Seydell-Greenwald et al., 2025)) as these may reveal regions with higher spatial overlap, given that language and ToM would be specialized to the same hemisphere, or if they are still distinct, help us better understand the importance of preserving modularity even in cases of unique cortical development.

The superior temporal lobe contains regions specialized for many different functions. In this paper, we show that language and ToM are functionally and spatially distinct and have distinct functional connectivity patterns to support each function within the STL. Our study aligns with prior work in adults and demonstrates distinct neural processors for language and ToM in for the first time in the developing human brain. We also show that connectivity is predictive of individualized activation and highlight the role of functional connectivity in shaping the functional responses of two communicative functions. Longitudinal analyses demonstrate that language and ToM do not disentangle over time. Overall, this work sheds light on the development of neural specialization and supports the idea that functional connectivity patterns are in place early in development and may scaffold the development of functionally specialized brain regions for uniquely human cognition.

## Acknowledgements

The authors are extremely grateful for the families who kindly supported this work. The authors would like to thank all current and past members of the Saygin Developmental Cognitive Neuroscience Lab for data collection.

## References

1. Astington, J. W., & Jenkins, J. M. (1999). A longitudinal study of the relation between language and theory-of-mind development. Developmental Psychology, 35, 1311–1320. 10.1037/0012-1649.35.5.1311

2. Baldwin, D. A., & Moses, L. J. (2001). Links between Social Understanding and Early Word Learning: Challenges to Current Accounts. Social Development, 10(3), 309–329. 10.1111/1467-9507.00168

3. Behzadi, Y., Restom, K., Liau, J., & Liu, T. T. (2007). A component based noise correction method (CompCor) for BOLD and perfusion based fMRI. NeuroImage, 37(1), 90–101. 10.1016/j.neuroimage.2007.04.042

4. Braga, R. M., DiNicola, L. M., Becker, H. C., & Buckner, R. L. (2020). Situating the left-lateralized language network in the broader organization of multiple specialized large-scale distributed networks. Journal of Neurophysiology, 124(5), 1415–1448. 10.1152/jn.00753.2019

5. Chai, L. R., Mattar, M. G., Blank, I. A., Fedorenko, E., & Bassett, D. S. (2016). Functional Network Dynamics of the Language System. Cerebral Cortex, 26(11), 4148–4159. 10.1093/cercor/bhw238

6. de Villiers, J. G. (2021). The Role(s) of Language in Theory of Mind. In M. Gilead & K. N. Ochsner (Eds.), The Neural Basis of Mentalizing (pp. 423–448). Springer International Publishing. 10.1007/978-3-030-51890-5_21

7. de Villiers, J. G., & de Villiers, P. A. (2014). The Role of Language in Theory of Mind Development. Topics in Language Disorders, 34(4), 313–328. 10.1097/TLD.0000000000000037

8. de Villiers, J. G., & Pyers, J. E. (2002). Complements to cognition: A longitudinal study of the relationship between complex syntax and false-belief-understanding. Cognitive Development, 17(1), 1037–1060. 10.1016/S0885-2014(02)00073-4

9. Deen, B., Koldewyn, K., Kanwisher, N., & Saxe, R. (2015). Functional Organization of Social Perception and Cognition in the Superior Temporal Sulcus. Cerebral Cortex, 25(11), 4596–4609. 10.1093/cercor/bhv111

10. Destrieux, C., Fischl, B., Dale, A., & Halgren, E. (2010). Automatic parcellation of human cortical gyri and sulci using standard anatomical nomenclature. NeuroImage, 53(1), 1–15. 10.1016/j.neuroimage.2010.06.010

11. Dronkers, N. F., Ludy, C. A., & Redfern, B. B. (1998). Pragmatics in the absence of verbal language: Descriptions of a severe aphasic and a language-deprived adult. Journal of Neurolinguistics, 11(1), 179–190. 10.1016/S0911-6044(98)00012-8

12. Dufour, N., Redcay, E., Young, L., Mavros, P. L., Moran, J. M., Triantafyllou, C., Gabrieli, J. D. E., & Saxe, R. (2013). Similar Brain Activation during False Belief Tasks in a Large Sample of Adults with and without Autism. PLOS ONE, 8(9), e75468. 10.1371/journal.pone.0075468

13. Ebert, S. (2020). Theory of mind, language, and reading: Developmental relations from early childhood to early adolescence. Journal of Experimental Child Psychology, 191, 104739. 10.1016/j.jecp.2019.104739

14. Enge, A., Friederici, A. D., & Skeide, M. A. (2020). A meta-analysis of fMRI studies of language comprehension in children. NeuroImage, 215, 116858. 10.1016/j.neuroimage.2020.116858

15. Fedorenko, E. (2021). The early origins and the growing popularity of the individual-subject analytic approach in human neuroscience. Current Opinion in Behavioral Sciences, 40, 105–112. 10.1016/j.cobeha.2021.02.023

16. Fedorenko, E., Hsieh, P.-J., Nieto-Castañón, A., Whitfield-Gabrieli, S., & Kanwisher, N. (2010). New Method for fMRI Investigations of Language: Defining ROIs Functionally in Individual Subjects. Journal of Neurophysiology, 104(2), 1177–1194. 10.1152/jn.00032.2010

17. Fedorenko, E., Ivanova, A. A., & Regev, T. I. (2024). The language network as a natural kind within the broader landscape of the human brain. Nature Reviews Neuroscience, 25(5), 289–312. 10.1038/s41583-024-00802-4

18. Freiwald, W. A. (2020). Social interaction networks in the primate brain. Current Opinion in Neurobiology, 65, 49–58. 10.1016/j.conb.2020.08.012

19. Garfield, J. L., Peterson, C. C., & Perry, T. (2001). Social Cognition, Language Acquisition and The Development of the Theory of Mind. Mind & Language, 16(5), 494–541. 10.1111/1468-0017.00180

20. Greve, D. N., & Fischl, B. (2009). Accurate and robust brain image alignment using boundary-based registration. NeuroImage, 48(1), 63–72. 10.1016/j.neuroimage.2009.06.060

21. Guan, Y., Keil, A., & Farrar, M. J. (2020). Electrophysiological dynamics of false belief understanding and complementation syntax in school-aged children: Oscillatory brain activity and event-related potentials. Journal of Experimental Child Psychology, 198, 104905. 10.1016/j.jecp.2020.104905

22. Gupta, M. D., Thakurta, R., & Basu, A. (2025). Relationship between Laterality and Theory of Mind among Typical Adults – A Systematic Literature Review. Acta Psychologica, 254, 104862. 10.1016/j.actpsy.2025.104862

23. Hale, C. M., & Tager-Flusberg, H. (2003). The Influence of Language on Theory of Mind: A Training Study. Developmental Science, 6(3), 346–359. 10.1111/1467-7687.00289

24. Hertrich, I., Dietrich, S., & Ackermann, H. (2020). The Margins of the Language Network in the Brain. Frontiers in Communication, 5, 519955. 10.3389/fcomm.2020.519955

25. Hiersche, K. J., Schettini, E., Li, J., & Saygin, Z. M. (2024). Functional dissociation of the language network and other cognition in early childhood. Human Brain Mapping, 45(9), e26757. 10.1002/hbm.26757

26. Ivanova, A., Zaidel, E., Salamon, N., Bookheimer, S., Uddin, L. Q., & de Bode, S. (2017). Intrinsic functional organization of putative language networks in the brain following left cerebral hemispherectomy. Brain Structure & Function, 222(8), 3795–3805. 10.1007/s00429-017-1434-y

27. Jacoby, N., Bruneau, E., Koster-Hale, J., & Saxe, R. (2016). Localizing Pain Matrix and Theory of Mind networks with both verbal and non-verbal stimuli. NeuroImage, 126, 39–48. 10.1016/j.neuroimage.2015.11.025

28. Johnson, M. H. (2011). Interactive Specialization: A domain-general framework for human functional brain development? Developmental Cognitive Neuroscience, 1(1), 7–21. 10.1016/j.dcn.2010.07.003

29. Kanwisher, N. (2010). Functional specificity in the human brain: A window into the functional architecture of the mind. Proceedings of the National Academy of Sciences, 107(25), 11163–11170. 10.1073/pnas.1005062107

30. Kraus, B. T., Perez, D., Ladwig, Z., Seitzman, B. A., Dworetsky, A., Petersen, S. E., & Gratton, C. (2021). Network variants are similar between task and rest states. NeuroImage, 229, 117743. 10.1016/j.neuroimage.2021.117743

31. Lenneberg, E. H. (1967). The Biological Foundations of Language. Hospital Practice, 2(12), 59–67. 10.1080/21548331.1967.11707799

32. Li, J., Hiersche, K. J., & Saygin, Z. M. (2024). Demystifying visual word form area visual and nonvisual response properties with precision fMRI. iScience, 27(12). 10.1016/j.isci.2024.111481

33. Malik-Moraleda, S., Ayyash, D., Gallée, J., Affourtit, J., Hoffmann, M., Mineroff, Z., Jouravlev, O., & Fedorenko, E. (2021). *The universal language network: A cross-linguistic investigation spanning 45 languages and 12 language families* [Preprint]. Neuroscience. 10.1101/2021.07.28.454040

34. Mars, R. B., Passingham, R. E., & Jbabdi, S. (2018). Connectivity Fingerprints: From Areal Descriptions to Abstract Spaces. Trends in Cognitive Sciences, 22(11), 1026–1037. 10.1016/j.tics.2018.08.009

35. Miller, C. A. (2006). Developmental Relationships Between Language and Theory of Mind. American Journal of Speech-Language Pathology, 15(2), 142–154. 10.1044/1058-0360(2006/014)

36. Molloy, M. F., Saygin, Z. M., & Osher, D. E. (2024). Predicting high-level visual areas in the absence of task fMRI. Scientific Reports, 14(1), 11376. 10.1038/s41598-024-62098-9

37. Ojemann, G. (1991). Cortical organization of language. The Journal of Neuroscience, 11(8), 2281–2287. 10.1523/JNEUROSCI.11-08-02281.1991

38. Oller, J. W., Oller, S. D., & Oller, S. N. (2012). Milestones: Normal Speech and Language Development Across the Lifespan. Plural Publishing.

39. Olulade, O. A., Seydell-Greenwald, A., Chambers, C. E., Turkeltaub, P. E., Dromerick, A. W., Berl, M. M., Gaillard, W. D., & Newport, E. L. (2020). The neural basis of language development: Changes in lateralization over age. Proceedings of the National Academy of Sciences, 117(38), 23477–23483. 10.1073/pnas.1905590117

40. Osher, D. E., Brissenden, J. A., & Somers, D. C. (2019). Predicting an individual’s dorsal attention network activity from functional connectivity fingerprints. Journal of Neurophysiology, 122(1), 232–240. 10.1152/jn.00174.2019

41. Osher, D. E., Saxe, R. R., Koldewyn, K., Gabrieli, J. D. E., Kanwisher, N., & Saygin, Z. M. (2016). Structural Connectivity Fingerprints Predict Cortical Selectivity for Multiple Visual Categories across Cortex. Cerebral Cortex, 26(4), 1668–1683. 10.1093/cercor/bhu303

42. Passingham, R. E., Stephan, K. E., & Kötter, R. (2002). The anatomical basis of functional localization in the cortex. Nature Reviews Neuroscience, 3(8), 606–616. 10.1038/nrn893

43. Paunov, A. M., Blank, I. A., & Fedorenko, E. (2019). Functionally distinct language and Theory of Mind networks are synchronized at rest and during language comprehension. Journal of Neurophysiology, 121(4), 1244–1265. 10.1152/jn.00619.2018

44. Paunov, A. M., Blank, I. A., Jouravlev, O., Mineroff, Z., Gallée, J., & Fedorenko, E. (2022). Differential Tracking of Linguistic vs. Mental State Content in Naturalistic Stimuli by Language and Theory of Mind (ToM) Brain Networks. Neurobiology of Language, 3(3), 413–440. 10.1162/nol_a_00071

45. Perner, J., Leekam, S. R., & Wimmer, H. (1987). Three-year-olds’ difficulty with false belief: The case for a conceptual deficit. British Journal of Developmental Psychology, 5(2), 125–137. 10.1111/j.2044-835X.1987.tb01048.x

46. Power, J. D., Barnes, K. A., Snyder, A. Z., Schlaggar, B. L., & Petersen, S. E. (2012). Spurious but systematic correlations in functional connectivity MRI networks arise from subject motion. Neuroimage, 59(3), 2142–2154. 10.1016/j.neuroimage.2011.10.018

47. Rajimehr, R., Firoozi, A., Rafipoor, H., Abbasi, N., & Duncan, J. (2022). Complementary hemispheric lateralization of language and social processing in the human brain. Cell Reports, 41(6), 111617. 10.1016/j.celrep.2022.111617

48. Rakoczy, H. (2022). Foundations of theory of mind and its development in early childhood. Nature Reviews Psychology, 1(4), 223–235. 10.1038/s44159-022-00037-z

49. Reuter, M., Rosas, H. D., & Fischl, B. (2010). Highly accurate inverse consistent registration: A robust approach. NeuroImage, 53(4), 1181–1196. 10.1016/j.neuroimage.2010.07.020

50. Richardson, H., Lisandrelli, G., Riobueno-Naylor, A., & Saxe, R. (2018). Development of the social brain from age three to twelve years. Nature Communications, 9(1), Article 1. 10.1038/s41467-018-03399-2

51. Richardson, H., & Saxe, R. (2020). Development of predictive responses in theory of mind brain regions. Developmental Science, 23(1), e12863. 10.1111/desc.12863

52. Saxe, R., Brett, M., & Kanwisher, N. (2006). Divide and conquer: A defense of functional localizers. NeuroImage, 30(4), 1088–1096. 10.1016/j.neuroimage.2005.12.062

53. Saygin, Z. M., Osher, D. E., Koldewyn, K., Reynolds, G., Gabrieli, J. D. E., & Saxe, R. R. (2012). Anatomical connectivity patterns predict face selectivity in the fusiform gyrus. Nature Neuroscience, 15(2), Article 2. 10.1038/nn.3001

54. Schettini, E., Hiersche, K. J., & Saygin, Z. M. (2023). Individual Variability in Performance Reflects Selectivity of the Multiple Demand Network among Children and Adults. Journal of Neuroscience, 43(11), 1940–1951. 10.1523/JNEUROSCI.1460-22.2023

55. Seydell-Greenwald, A., Vladyko, N., Chambers, C. E., Gaillard, W. D., Landau, B., & Newport, E. L. (2025). Right-Lateralization of the Visual Word Form Area after Left-Hemisphere Perinatal Stroke. Journal of Neuroscience, 45(10). 10.1523/JNEUROSCI.0924-24.2024

56. Shain, C., Paunov, A., Chen, X., Lipkin, B., & Fedorenko, E. (2022). No evidence of theory of mind reasoning in the human language network. *Cerebral Cortex*, bhac505. 10.1093/cercor/bhac505

57. Sharp, H., & Hillenbrand, K. (2008). Speech and Language Development and Disorders in Children—ClinicalKey. 55(5), 1159–1173.

58. Skeide, M. A., & Friederici, A. D. (2016). The ontogeny of the cortical language network. Nature Reviews Neuroscience, 17(5), Article 5. 10.1038/nrn.2016.23

59. Slade, L., & Ruffman, T. (2005). How language does (and does not) relate to theory of mind: A longitudinal study of syntax, semantics, working memory and false belief. British Journal of Developmental Psychology, 23(1), 117–141. 10.1348/026151004X21332

60. Smith, S. M., Fox, P. T., Miller, K. L., Glahn, D. C., Fox, P. M., Mackay, C. E., Filippini, N., Watkins, K. E., Toro, R., Laird, A. R., & Beckmann, C. F. (2009). Correspondence of the brain’s functional architecture during activation and rest. Proceedings of the National Academy of Sciences, 106(31), 13040–13045. 10.1073/pnas.0905267106

61. Tavor, I., Jones, O. P., Mars, R. B., Smith, S. M., Behrens, T. E., & Jbabdi, S. (2016). Task-free MRI predicts individual differences in brain activity during task performance. Science, 352(6282), 216–220. 10.1126/science.aad8127

62. Tik, N., Gal, S., Madar, A., Ben-David, T., Bernstein-Eliav, M., & Tavor, I. (2023). Generalizing prediction of task-evoked brain activity across datasets and populations. NeuroImage, 276, 120213. 10.1016/j.neuroimage.2023.120213

63. Tobyne, S. M., Somers, D. C., Brissenden, J. A., Michalka, S. W., Noyce, A. L., & Osher, D. E. (2018). Prediction of individualized task activation in sensory modality-selective frontal cortex with ‘connectome fingerprinting.’ NeuroImage, 183, 173–185. 10.1016/j.neuroimage.2018.08.007

64. Tylén, K., Weed, E., Wallentin, M., Roepstorff, A., & Frith, C. D. (2010). Language as a Tool for Interacting Minds. Mind & Language, 25(1), 3–29. 10.1111/j.1468-0017.2009.01379.x

65. Varley, R., Siegal, M., & Want, S. C. (2001). Severe Impairment in Grammar Does Not Preclude Theory of Mind. Neurocase, 7(6), 489–493. 10.1093/neucas/7.6.489

66. Weiss-Croft, L., & Baldeweg, T. (2015). Maturation of language networks in children: A systematic review of 22years of functional MRI | Elsevier Enhanced Reader. 123, 269–281. 10.1016/j.neuroimage.2015.07.046

67. Willems, R. M., Benn, Y., Hagoort, P., Toni, I., & Varley, R. (2011). Communicating without a functioning language system: Implications for the role of language in mentalizing. Neuropsychologia, 49(11), 3130–3135. 10.1016/j.neuropsychologia.2011.07.023

