## Supplemental Table and Figures for "Functional Dissociation in the Developing Superior Temporal Lobe: Language and Theory of Mind"

Supplemental Materials:

**Supplemental Tables:**

- Table S1: Selectivity of Language and ToM fROIs for adults and children.
- Table S2: Age and development related selectivity changes in fROIs to non-preferred categories in children.
- Table S3: Overlap of maximally sensitive regions across thresholds for adults and children.
- Table S4: Age and development changes of overlap in children.

**Supplemental Figures:**

- Figure S1: Group level laterality analysis for language and ToM in adults and children.
- Figure S2: Top 10 regions with the most disparate beta values across language and ToM CF models in adults.
- Figure S3: Beta weight values for regions with greatest differences between child and adult models

| Adults | | | | Children | | |
|---|---|---|---|---|---|---|
| **Language Selectivity** | | | | **Language Selectivity** | | |
| *fROI* | *t stat (dof)* | *p value* | *d* | *t stat (dof)* | *p value* | *D* |
| l-lang | 14.75 (28) | 4.9614e-15** | 2.66 | 8.17 (55) | 2.2843e-11** | 1.08 |
| r-lang | 10.44 (28) | 1.8319e-11** | 1.89 | 7.14 (55) | 1.1312e-09** | 0.94 |
| l-ToM | 5.74 (27) | 2.1085e-06** | 1.05 | 3.09 (54) | 0.0016007** | 0.41 |
| r-ToM | 1.79 (28) | 0.042486** | 0.32 | 0.25 (53) | 0.40283 | 0.03 |
| l-langAG | 4.04 (28) | 0.000191** | 0.73 | 3.86 (54) | 0.00015169** | 0.51 |
| r-langAG | 2.45 (27) | 0.01046** | 0.45 | 4.10 (50) | 7.5135e-05** | 0.57 |
| r-tomSTS | 4.96 (28) | 1.5427e-05** | 0.9 | 2.89 (53) | 0.0028224** | 0.39 |
| **ToM Selectivity** | | | | **ToM Selectivity** | | |
| *fROI* | *t stat (dof)* | *p value* | *d* | *t stat (dof)* | *p value* | *d* |
| l-lang | 6.30 (28) | 4.0455e-07** | 1.14 | 4.81 (54) | 6.1667e-06** | 0.64 |
| r-lang | 5.33 (28) | 5.592e-06** | 0.96 | 5.83 (54) | 1.5767e-07** | 0.78 |
| l-ToM | 7.58 (28) | 1.485e-08** | 1.37 | 6.87 (53) | 3.6365e-09** | 0.92 |
| r-ToM | 7.04 (28) | 5.8496e-08** | 1.27 | 8.06 (53) | 4.5344e-11** | 1.08 |
| l-langAG | 4.30 (28) | 9.3522e-05** | 0.78 | 2.70 (51) | 0.0046908** | 0.37 |
| r-langAG | 0.18 (27) | 0.42844 | 0.03 | 3.48 (50) | 0.00052179** | 0.48 |
| r-tomSTS | 4.35 (28) | 8.1914e-05** | 0.79 | 2.18 (53) | 0.016687** | 0.29 |

**Table S1:** Results of one-tailed t-tests for testing significance of selectivity to language and ToM in each fROI.

| Selectivity changes with age | | | | |
|---|---|---|---|---|
| | Language | | ToM | |
| | r | p | r | p |
| l-lang | 0.434 | 0.00093** | 0.156 | 0.261 |
| r-lang | 0.021 | 0.877 | 0.050 | 0.717 |
| l-ToM | -0.011 | 0.937 | 0.027 | 0.849 |
| r-ToM | -0.151 | 0.281 | -0.009 | 0.949 |
| l-langAG | -0.058 | 0.676 | -0.153 | 0.285 |
| r-langAG | -0.107 | 0.458 | -0.106 | 0.463 |
| r-tomSTS | -0.032 | 0.819 | -0.115 | 0.413 |

| Selectivity comparison across TPs | | | | | | |
|---|---|---|---|---|---|---|
| | Language | | | ToM | | |
| | *t(df)* | *p* | *d* | *t(df)* | *p* | *d* |
| l-lang | -1.78 (19) | 0.090 | -0.529 | -0.70 (19) | 0.492 | -0.188 |
| r-lang | -1.84 (18) | 0.082 | -0.507 | -0.16 (19) | 0.873 | -0.054 |
| l-ToM | -1.94 (15) | 0.072 | -0.537 | 1.53 (16) | 0.146 | 0.094 |
| r-ToM | -1.23 (16) | 0.237 | -0.246 | 0.71 (18) | 0.484 | 0.164 |
| l-langAG | -1.53 (18) | 0.144 | -0.442 | -0.03 (18) | 0.975 | -0.042 |
| r-langAG | -2.48 (17) | 0.024* | -1.007 | -0.82 (17) | 0.423 | -0.193 |
| r-tomSTS | -1.30 (16) | 0.213 | -0.236 | 1.64 (16) | 0.121 | 0.493 |
| r-tomSTS | -1.30 (16) | 0.213 | -0.236 | 1.64 (16) | 0.121 | 0.493 |

**Table S2:** Top: Results of partial correlation between age and selectivity (controlling for framewise displacement) in each fROI. Bottom: Results of paired samples t-test to compare selectivity across timepoints to language and ToM in each fROI. *p<0.05, not surviving multiple comparison correction; **p<0.05 after Bonferroni-Holm correction for 7 comparisons.

| Overlap of maximally sensitive regions | | | | |
|---|---|---|---|---|
| | Adults | | Children | |
| | LH STL | RH STL | LH STL | RH STL |
| *Threshold* | *mean (sd)* | *mean (sd)* | *mean (sd)* | *mean (sd)* |
| 1% | 0.048 (0.020) | 0.090 (0.026) | 0.014 (0.005) | 0.030 (0.008) |
| 2% | 0.057 (0.021) | 0.096 (0.025) | 0.019 (0.005) | 0.037 (0.009) |
| 3% | 0.074 (0.021) | 0.112 (0.025) | 0.029 (0.006) | 0.050 (0.011) |
| 4% | 0.087 (0.021) | 0.125 (0.025) | 0.038 (0.006) | 0.065 (0.013) |
| 5% | 0.101 (0.021) | 0.140 (0.025) | 0.048 (0.007) | 0.075 (0.014) |
| 10% | 0.173 (0.023) | 0.201 (0.022) | 0.095 (0.010) | 0.123 (0.015) |
| 20% | 0.284 (0.020) | 0.303 (0.018) | 0.193 (0.012) | 0.212 (0.014) |
| 30% | 0.383 (0.017) | 0.390 (0.015) | 0.297 (0.012) | 0.310 (0.012) |

**Table S3:** Mean (sd) overlap of language and ToM within hemisphere at all thresholds.

| Correlation of overlap and age | | | | | | |
|---|---|---|---|---|---|---|
| | LH STL | | RH STL | | LH Lang - RH ToM | |
| *Threshold* | *r* | *p* | *r* | *p* | *r* | *p* |
| 1% | 0.09 | 0.49 | -0.087 | 0.528 | 0.20 | 0.13 |
| 2% | 0.04 | 0.80 | -0.046 | 0.735 | 0.16 | 0.23 |
| 3% | -0.08 | 0.58 | -0.082 | 0.549 | 0.23 | 0.08 |
| 4% | -0.12 | 0.37 | -0.090 | 0.510 | 0.21 | 0.11 |
| 5% | -0.14 | 0.32 | -0.103 | 0.450 | 0.19 | 0.16 |
| 10% | -0.08 | 0.54 | -0.043 | 0.755 | 0.05 | 0.69 |
| 20% | 0.02 | 0.88 | 0.048 | 0.728 | 0.13 | 0.32 |
| 30% | 0.09 | 0.53 | 0.122 | 0.370 | 0.16 | 0.25 |

| Overlap Comparison Across TPs | | | | | | | | | |
|---|---|---|---|---|---|---|---|---|---|
| *Threshold* | LH STL | | | RH STL | | | LH Lang - RH ToM | | |
| | *t(df)* | *p* | *d* | *t(df)* | *p* | *d* | *t(df)* | *p* | *d* |
| 1% | -1.33 (19) | 0.20 | -0.4 | 2.30 (17) | 0.034$^{+}$ | 0.77 | 0.00 (19) | 1.000 | 0.000 |
| 2% | -1.22 (19) | 0.24 | -0.36 | 2.12 (19) | 0.048$^{+}$ | 0.67 | -0.04 (19) | 0.972 | -0.011 |
| 3% | -0.77 (19) | 0.45 | -0.21 | 1.91 (19) | 0.071 | 0.6 | 0.57 (19) | 0.575 | 0.177 |
| 4% | -0.55 (19) | 0.59 | -0.13 | 1.65 (19) | 0.116 | 0.53 | 0.42 (19) | 0.679 | 0.122 |
| 5% | -0.38 (19) | 0.71 | -0.09 | 1.73 (19) | 0.101 | 0.54 | 0.27 (19) | 0.791 | 0.076 |
| 10% | -0.37 (19) | 0.72 | -0.1 | 1.47 (19) | 0.157 | 0.48 | 0.07 (19) | 0.947 | 0.019 |
| 20% | -0.20 (19) | 0.84 | -0.06 | 1.46 (19) | 0.160 | 0.45 | -0.18 (19) | 0.862 | -0.054 |
| 30% | 0.02 (19) | 0.99 | 0 | 1.26 (19) | 0.221 | 0.34 | -0.55 (19) | 0.590 | -0.187 |

**Table S4:** Top: Results of correlation between age and language-to-ToM overlap within the same hemisphere and the spatial overlap of LH language with RH ToM at each threshold. Bottom: Results of paired-samples t-tests to compare overlap across timepoints. $^{+}p<0.05$, not surviving multiple comparison correction for 8 thresholds.

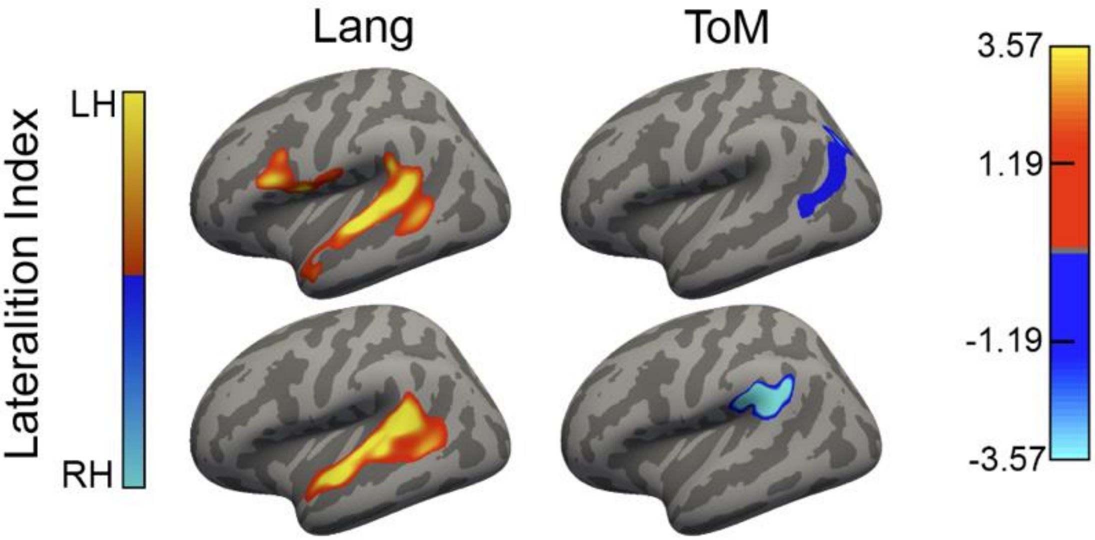


**Figure S1:** Group-level lateralization of activation for language and theory of mind in adults and children. Lateralization maps are shown on the inflated cortical surface (fsaverage_sym). The top row displays results for adults, and the bottom row displays results for children.

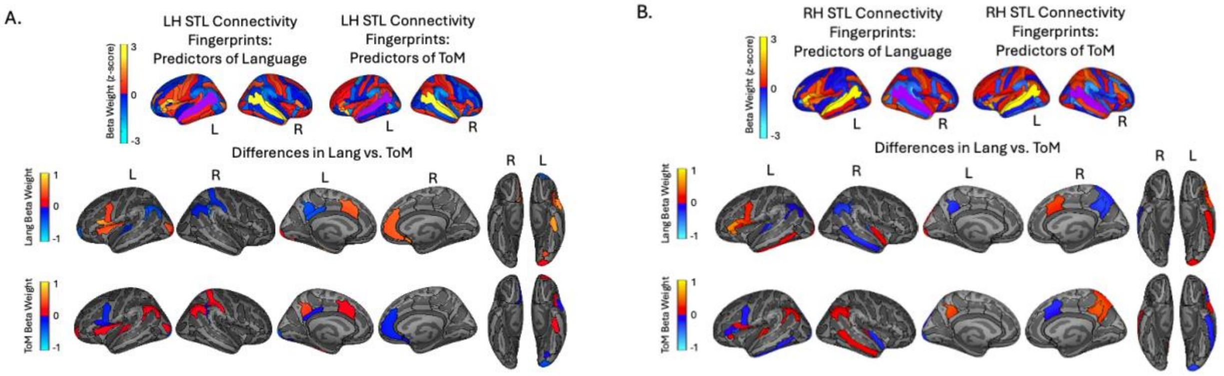


**Figure S2:** Beta value for the adult language and ToM models. Z-score of beta weights for language and ToM models for A. LH and B. RH. Top row shows beta weights of predictors on the lateral surface for illustrative purposes. The region being predicted is depicted in purple. Below, beta weights for regions that are most different when predicting language and ToM (10 most positive and 10 most negative t-values after paired-sample t-tests across language and ToM models) for LH and RH separately. Middle row depicts beta value in the model predicting language function, bottom row depicts beta value for the model predicting ToM function.

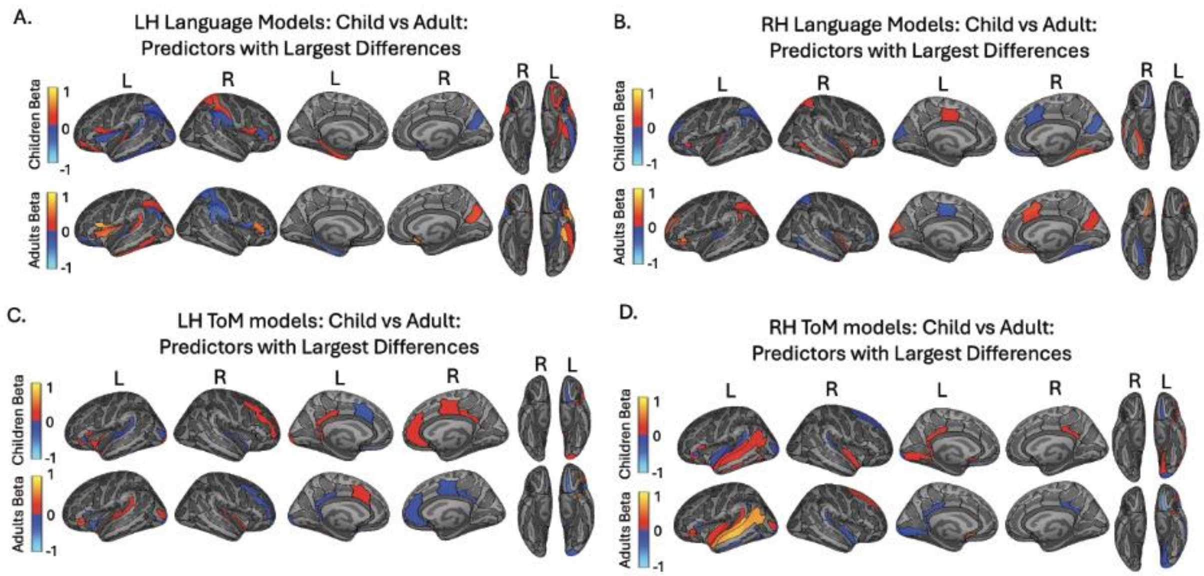


**Figure S3** Beta weight values for regions with greatest differences between child and adult models (top 10 positive and top 10 negative t-statistics for adult vs. child comparison). A. LH language model; B. RH language models; C. LH ToM models; D. RH ToM models. Top row depicts beta value in adult models, bottom row depicts beta value for the child models.
